# Photoluminescence brightening of single-walled carbon nanotubes through conjugation with graphene quantum dots

**DOI:** 10.1101/2023.02.28.528463

**Authors:** Sayyed Hashem Sajjadi, Shang-Jung Wu, Melania Reggente, Niloufar Sharif, Ardemis A. Boghossian

## Abstract

Spanning the tissue transparency window, the near-infrared (NIR) photoluminescence (PL) of single-walled carbon nanotubes (SWCNTs) can optically penetrate biological tissue for deep-tissue imaging and optical sensing. SWCNTs are often functionalized with single-stranded DNA (ssDNA) to yield biocompatible, responsive, and selective sensors. However, the low brightness of these ssDNA-wrapped SWCNTs sensors restricts the depth at which such sensors can be implanted in the tissue. This work demonstrates the PL enhancement of ssDNA-wrapped SWCNTs by incorporating biocompatible graphene quantum dots (GQDs). Two kinds of GQDs, pristine (PGQDs) and nitrogen-doped (NGQDs), were fabricated and characterized. Thermodynamically, both GQDs were shown to significantly increase the fluorescence efficiency of ssDNA-SWCNTs with the same degree of PL enhancement after 3 h. Furthermore, a correlation between the diameter of the SWCNTs and the PL enhancement factor was found; the larger the SWCNT diameter, the higher the PL increase upon adding GQDs. For instance, a 30-fold enhancement was achieved for (8,6) chirality while it was only 2-fold for the (6,5) chirality. Our experiments showed that adding GQDs increases the surface coverage of SWCNTs suspended by ssDNA, limiting water molecules’ access to the nanotube surface, thus increasing the fluorescence efficiency. Kinetically, NGQDs brightened SWCNTs much faster than PGQDs. The PL intensity reached a plateau in 2 min following the addition of NGQDs, while it was still increasing even after 1 h upon the addition of PGQDs. We show that NGQDs can act as reducing agents to decrease the amount of dissolved oxygen, which quenches the SWCNTs PL. This advancement provides a promising tool for engineering the brightness of NIR sensors for biomedical applications such as single-molecule imaging of individual SWCNTs using NIR confocal microscopy and deep tissue sensing.

## Introduction

Biological fluids and tissues are autofluorescent and absorb light below 950 nm. Water forms the basis of biofluids and shows strong absorption above 1350 nm. Therefore, biotransparent fluorophores that emit light in the second near-infrared (NIR) window (950-1350 nm) are advantageous for in-vivo and in-vitro imaging and optical sensing. Their emissions can penetrate deep into tissues without being absorbed or scattered, facilitating accurate and detailed imaging and sensing.^1^ Amongst these sensors, cylindrically rolled graphene sheets called single-walled carbon nanotubes (SWCNTs) show distinct intrinsic non-bleaching NIR photoluminescence (PL).^2^ For optical sensors based on the NIR PL of SWCNTs, specific attributes such as quantum yield (QY) and emission peak position can be modulated via surface functionalization.

The functionalization of SWCNTs is achieved in a covalent or non-covalent manner. Covalent functionalization may lead to PL brightening of SWCNTs.^3^ However, this approach not only destroys the extended π-network of SWCNTs^4^ but also requires expertise in organic chemistry and is expensive and time-consuming. On the other hand, the more straightforward approach of non-covalent functionalization enables researchers to prepare bright, sensitive, and selective SWCNT sensors faster and without needing particular expertise. Different wrappings, such as surfactants, polymers, and biomolecules (DNAs and proteins), have been used for the non-covalent functionalization and suspension of SWCNTs in solutions.^4^ Single-stranded DNA (ssDNA) is one of the extensively studied non-covalent surface wrappings. Due to their high colloidal stability in aqueous solutions and their specificity to analytes, SWCNTs functionalized with ssDNA are valuable for optical sensing applications.^5–12^ However, these complexes suffer from lower QYs (0.1–1%) compared with most conventional fluorophores, which reduces the penetration depth of these sensors.^6,13^

Several strategies have been proposed for enhancing the QY of SWCNTs in aqueous solutions, including directed evolution of DNA wrapping,^6^ addition of reducing agents,^14–16^ incorporation of defect quantum states,^17^ admixture with electrolytes,^18,19^ brightening by a dye through resonance energy transfer,^20^ and the use of plasmonic nanoparticles.^13,20^ The directed evolution method has been implemented to screen the ssDNA wrapping libraries, leading to a bright ssDNA-SWCNT conjugate.^6^ It is a novel approach that can increase the QY of ssDNA-SWCNT by 56% following two evolution cycles. A further approach involves using reductants like Trolox and dithiothreitol (DTT) to enhance the fluorescence of SWCNTs suspended by ssDNA or sodium dodecyl sulfate (SDS).^14,15^ Despite a greater than 10-fold QY enhancement, increased cytotoxicity and reduced sensor sensitivity limit the utilization of these reductant brightening agents. Recently, we reported plasmon-enhanced fluorescence using plasmonic nanoparticles,^13^ demonstrating that silver nanotriangles could increase the fluorescence intensity of ssDNA-SWCNT by up to 240%.

The literature also shows that SWCNTs can act as quenching agents when combined with inorganic quantum dots (QDs) or organic fluorophores due to the electron or energy transfer from the fluorophores to SWCNTs.^21^ Recently, it was demonstrated that the PL of graphene QDs (GQDs) is quenched in the presence of SWCNTs.^22^ Das et al. concluded that the charge transfer from GQDs to semiconducting SWCNTs is responsible for the static and dynamic quenching of GQDs due to a specific band alignment at the GQD-SWCNT interface. GQDs, single or few-layer zero-dimensional graphene sheets with diameter sizes of <20 nm, are fluorescent members of the nanocarbon family, standing for a kind of QDs with unique properties associated with both graphene and inorganic QDs.^23,24^ Compared to other fluorescent nanomaterials (e.g., organic dyes, inorganic QDs, and plasmonic nanoparticles), biocompatible GQDs are interesting materials for designing optoelectronic devices, sensors, bioimaging, photocatalysis, and photoelectrocatalysis due to their low cost, PL stability, and robust chemical inertness.^25–27^

In this work, we examine the effect of GQDs on the fluorescence of SWCNTs suspended by ssDNA for the first time. While SWCNTs are reported to quench the fluorescence of GQDs, investigating the change in the fluorescence of SWCNTs when conjugated with GQDs is lacking. Jeong et al.^28^ recently demonstrated that GQDs could interact with ssDNA strands depending on their oxidation level; ssDNA was adsorbed on the surface of GQDs with no or low oxidation, while no adsorption occurred in GQDs with medium and high oxidation levels. To avoid the interactions between GQDs and ssDNA, we synthesized three different types of medium-oxidation GQDs, compared their influence on the QY of ssDNA-SWCNTs of different chiralities, and explored the related mechanisms. The application of such conjugation was examined through NIR confocal microscopy and deep tissue fluorescence experiments.

## Materials and Methods

### Chemicals

All ssDNA oligomers were purchased from Microsynth (Microsynth AG, Switzerland). Super Purified HiPCO-SWCNTs were purchased from NanoIntegris (Lot. No. HS37-007, NanoIntegris Technologies, Canada). Oxygen (99.999%) and nitrogen (99.999%) gases were purchased from Carbagas (Carbagas AG, Switzerland). All other Chemicals were obtained from Sigma-Aldrich unless otherwise specified.

### Preparation nanomaterials

#### Fabrication of GQDs

Pristine graphene quantum dots (PGQDs) were prepared using the direct pyrolysis of citric acid (CA) method as previously described (**Fig. 1a**).^29^ In this process, 2g of CA was heated in a 5 mL beaker to 200°C. In 5 min, the CA was liquated. The color of the liquid changed from clear to pale yellow and then orange over the course of 30 minutes, indicating the formation of GQDs. For preparing PGQD solution, the obtained orange solution (A) was neutralized to pH 7.0 with NaOH. Next, PGQDs were reduced by hydrazine achieving nitrogen-doped GQDs (NGQDs, **Fig. 1**a).^29^ After neutralizing solution A, 2 ml of hydrazine-hydrate was added, and the mixture was stirred at 65 °C for 24 h. As a control, another kind of nitrogen-doped GQDs (GGQDs) were synthesized through microwave irradiation of glucosamine-HCl aqueous solution (**Fig. S1**).^30^ A 0.14 M aqueous solution of glucosamine-HCl was placed in a domestic microwave (Fust, 900 W) for 40 min at 50% power.

**Fig. 1.**
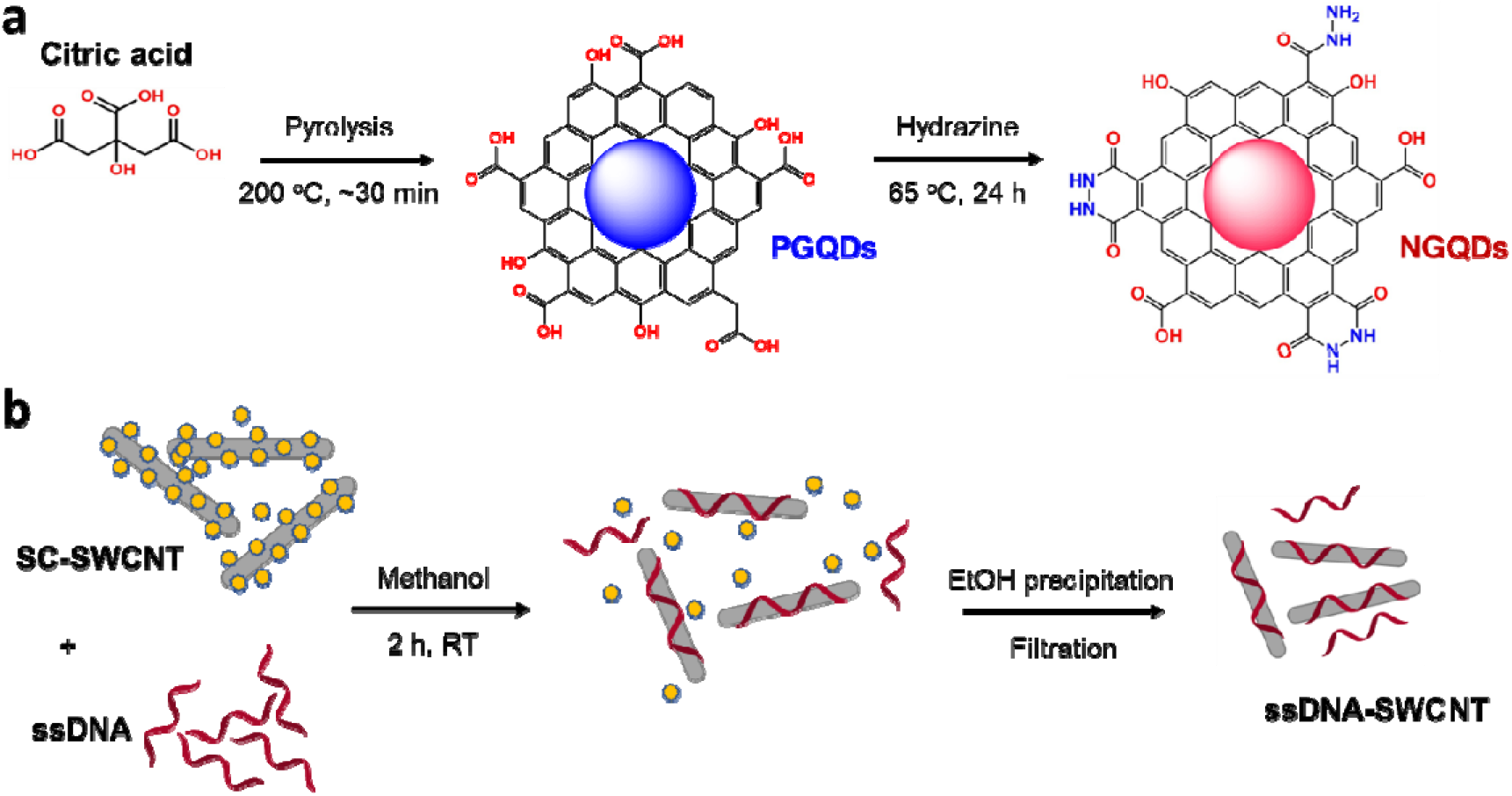
Preparation of graphene quantum dots (GQDs) and ssDNA-SWCNT samples. (a) Synthesis of pristine GQDs (PGQDs) through pyrolysis of citric acid and nitrogen-doped GQDs (NGQDs) through hydrazine treatment of PGQDs. (b) Preparation of ssDNA-SWCNT complexes via a surfactant exchange method in methanol.

The aqueous solutions of GQDs were purified utilizing dialysis kits and Amicon ultra-filtration devices (Amicon Ultra-15, Merck) with 1 and 3 kDa membranes, respectively. Before use, the dialysis kit and the filtration device were rinsed with DI water following the manufacturer’s recommendations. Finally, GQDs were precipitated using cold ethanol followed by centrifugation, dried, and kept in a vacuum desiccator.

#### Preparation of ssDNA-SWCNT solution

A previously reported method was used to prepare ssDNA-suspended SWCNTs.^6^ Briefly, wrapping exchange between surfactant-suspended SWCNTs and ssDNA was performed in the presence of methanol, which was added to increase the critical micelle concentration (CMC) of the surfactant. This study used sodium cholate (SC) as the dispersant for SWCNTs. 40 mg of SWCNTs were suspended in 40 mL of 2% (w/w) SC in DI water. The suspension was homogenized for 20 min at 5000 rpm (PT 1300D, Polytron) and sonicated for 1 h using a probe-tip ultrasonicator (1/4 in. tip, Q700 Sonicator, Qsonica) at 10% amplitude in a 4 °C water bath. Finally, the SWCNT suspension was centrifuged at 30000 rpm (164 000 x g) for 4 h at 25 °C (Optima XPN-80, Beckman Coulter) to remove any remaining nanotube aggregates.

To exchange the surfactant with ssDNA, 100 μL of SC-SWCNTs (108 mg/L in 2 wt% SC in ddH_2_O) were mixed with 100 μL of ssDNA solution (100 μM in DI water). DNA concentrations were measured and adjusted based on absorbance measurements (Nanodrop 2000, Thermo Scientific). Next, 300 μL of methanol (VWR Chemicals) was added to the ssDNA and SC-SWCNT mixture to obtain a final solvent percentage of 60% (v/v). The solution was vortexed briefly and subsequently incubated for 2 h at room temperature. Following the incubation, impurities (such as remaining catalyst particles, surfactant, and methanol) and unbound ssDNA were removed from the ssDNA-SWCNT suspensions. The obtained solutions were purified using Amicon centrifugal devices (Amicon Ultra-0.5,100 kDa membrane, Merck). The solution of ssDNA-SWCNTs was added to the filtration device and rinsed eight times (at 4,000 × g for 2 min) with 0.45 mL aliquots of DI water. The rinsed suspension was collected from the filtration device and centrifuged for a minimum of 1 h at 21,130 × g and 4 °C to remove any additional SWCNT aggregates that may have formed during the rinsing process.

#### Preparation of GQD-conjugated ssDNA-SWCNTs

GQD-conjugated ssDNA-SWCNTs were prepared by mixing 48 μL of a 2 mg/L of ssDNA-SWCNT solution (as estimated from the absorption spectrum, ε632nm = 36 mL mg^−1^cm^−1^) with 2 μL of different GQD solutions (with the final concentration of 0.1 mg/ml). The mixtures were incubated for at least 3 h before analyses unless otherwise specified.

#### X-ray photoelectron spectroscopy

X-ray photoelectron spectroscopy (XPS) measurements were carried out on a PHI VersaProbe II scanning XPS microprobe (Physical Instruments AG, Germany) to determine the chemical composition of GQDs. Data analysis and peak deconvolutions were performed on Casa XPS software.

#### Transmission electron microscopy

Transmission electron microscopy (TEM) and energy dispersive X-ray (EDX) analyses were performed on an FEI Tecnai Osiris at an acceleration voltage of 200 kV. This microscope has a high-brightness X-FEG gun and silicon drift Super-X EDX detectors. High-angle annular dark-field (HAADF) images were acquired in scanning TEM (STEM) mode. TEM samples were prepared by drop-casting solutions of nanoparticles onto Cu grids. TEM images were processed using DigitalMicrograph software (Gatan, USA).

#### Atomic Force microscopy

Atomic Force microscopy (AFM) was used to characterize the morphology of the nanoparticles and performed on a commercial microscope (Cypher, Asylum Research) equipped with a Si cantilever (AC240TS-R3, Asylum Research). Topography and amplitude images were acquired in standard tapping mode. Nanoparticle (GQDs, ssDNA-SWCNTs, or ssDNA-SWCNTs + GQDs) suspensions (20 μL) were drop-casted onto freshly cleaved mica substrates and dried at room temperature. Images were analyzed in the AFM data analysis software (Gwyddion 2.52).

#### UV-Vis absorbance and photoluminescence spectroscopy

Absorbance spectra of different nanoparticle solutions were recorded using a UV-Vis-NIR scanning spectrometer (Shimadzu 3600 Plus) with a 50 µL quartz cuvette (Suprasil quartz, path length 3 mm, Hellma). UV-Vis PL spectra and Vis PL emission-excitation (PLE) maps were acquired using a JASCO FP-6500 composed of two monochromators for excitation and emission, a 150 Watt Xe lamp with a shielded lamp house, and a photomultiplier as a light detector.

#### Cyclic voltammetry

Cyclic voltammetry (CV) measurements were performed in a conventional three-electrode electrochemical cell using a Pt wire and a standard Ag/AgCl (in saturated KCl solution) as counter and reference electrodes, respectively. A glassy carbon served as the working electrode. The setup was connected to a PalmSens4 potentiostat (PalmSens BV) equipped with PSTrace software. To measure the electroactivity of the as-synthesized GQDs, a 100 mM NaCl buffer (as blank) supplemented with 1 mg/mL of different types of GQDs were employed as electrolyte solutions, and cyclic voltammograms were recorded at different scan rates (5 mV/s, 10 mV/s, and 20 mV/s).

#### NIR photoluminescence spectroscopy

NIR-II PL spectra of the samples were collected using a custom-built NIR optical setup on an inverted Nikon Eclipse Ti-E microscope (Nikon AG Instruments), as described previously.^6^ Sample preparation was carried out in 384-well plates (UV-star, Greiner). The setup consists of a supercontinuum laser source and a tunable band-pass filter unit (SuperK Extreme EXR-15 and SuperK Varia, NKT Photonics) that operates between 400 and 830 nm at a pulse frequency of 80 MHz. Light emitted from the sample is focused onto an SCT-320 spectrometer (Princeton Instruments) and then redirected into an InGaAs NIR camera (NIRvana 640 ST, Princeton Instruments). A grating of 75 mm^−1^ was used to collect the nanotube emission between 850 – 1350 nm. Measurements were recorded with LightField (Princeton Instruments, USA) and a custom-built LabView (National Instruments) software. An exposure time of 10 s and laser excitation with a bandwidth of 10 nm and relative power of 100% was used for all measurements unless stated otherwise. Spectral fitting was performed using a custom Python program. NIR PLE maps were acquired between 500 nm and 800 nm with a 5 nm step, and the results were analyzed using a custom MATLAB code (MATLAB R2018b, MathWorks).

Deep-tissue NIR spectra were collected on the same setup. Ham tissue slices of different thicknesses (0 mm – 2.18 mm) were inserted between the ssDNA-SWCNTs (2 mg/l) (with or without NGQDs) and the laser excitation source (730 nm).

#### Gas purging NIR photoluminescence spectroscopy

Gas experiments were performed on an inverted Axiovert 200M microscope (Zeiss). The setup consisted of a supercontinuum laser (NKT Photonics) that operated between 400 and 830 nm. Light emitted from the sample was focused onto an iHR320 spectrometer (Horiba Scientific) and then redirected into a liquid nitrogen-cooled InGaAs NIR camera (Symphony II, Horiba Scientific). A grating of 150 mm^−1^ was used to collect the nanotube emission between 900 – 1400 nm. 400 μL of ssDNA-SWCNTs (5 mg/L) solution was poured into a transparent flat-bottom glass vial. Nitrogen gas (N_2_) with a flow rate of 0.05 L/min was purged inside the solution for 1 h before acquiring the initial spectrum. After purging N_2_, oxygen gas (O_2_, 0.1 L/min) was purged for another 1 h, and the spectrum was collected. To investigate the effect of additives on the O_2_ saturated samples, either GQDs or DTT with the final concentrations of 1 mg/mL or 1 mM, respectively, were added to the container following O_2_ purging for 15 min, and then the spectra were collected. To avoid the effect of bubbling on the PL intensities, the inlet gas pipe was out of solution (but in the vial) when the PL spectra were recorded. Gas flow rates were controlled by a SmartTrak@ series / Pilot Module mass flowmeter (Vögtlin Instruments GmbH, Switzerland), and the minimum possible flow rates were chosen. The vial was sealed with a cap (containing 2 tiny holes for the gas inlet and outlet) to minimize the evaporation of the solution during gas purging. It is worth noting that the concentration (and the volume) of the ssDNA-SWCNTs solution was measured after gas purging, and it was quite the same as the initial concentration (∼ 400 μL, 5 mg/L), showing that the evaporation did not affect the PL intensities.

#### Wide-field and confocal fluorescence microscopy

Nanotubes were coated onto poly-L-lysine coated glass Petri dishes by incubating 50 μL of (GT)_20_-SWCNT solution (2 mg/L) for 30 min, followed by washing with DI water.

As described previously, single-molecule images of SWCNT were taken using a custom-built NIR optical setup on an inverted Nikon Eclipse Ti-E microscope (Nikon AG Instruments).^1^ A 640□nm continuous wave (CW) laser light source (Triline Laser Bank, Cairn Research) was coupled to the microscope body by an optical fiber (FT1500 EMT, 0.39 NA). The laser light passed through a spinning-disc confocal unit (X-light, CrestOptics) that included discs of spinning arrays with pinholes (ø 60□µm) and lenses coated with a NIR anti-reflection layer (transmittance >60% in the wavelength range from 0.7 to 1.2□µm). A dichroic mirror splits the excitation and emission beams. Samples mounted in the XYZ-translational stage were all illuminated through a TIRF Apo 100□×□1.49 NA oil immersion objective (Nikon Instruments) and a 1.5x tube lens. The fluorescence signal was collected in the epi-direction through a 980 ± 15□nm band-pass (Chroma Technology) or a 980□nm long-pass (BLP01-980R-25, Semrock) filter by a cooled indium gallium arsenide (InGaAs) camera (NIRvana 640 ST, Princeton Instruments).

Wide-field visible fluorescence images of samples were collected by the same microscope setup (Nikon Ti-E) equipped with a Shamrock 303i spectrometer (Andor) coupled to EMCCD visible camera (iXon Ultra 888, Andor). A 370 nm LED was used as the excitation source.

NIR and visible fluorescence images were acquired using the Nikon NIS-Elements software (Nikon Instruments) and were analyzed using a custom Matlab code (Matlab R2018b, Mathworks).

## Results and Discussion

We characterized the structure of GQDs by TEM and AFM. **Fig. 2a** (top) shows the TEM image of PGQDs of round shape and average diameter size of 13.1 nm according to the size distribution plot in **Fig. 2a** (bottom). Although NGQDs were obtained through the reaction of PGQDs and hydrazine, their sizes were smaller than half of those for PGQDs (≈ 4.8 nm, **Fig. 2b**), implying the shrinkage of PGQDs to NGQDs during the heating process. AFM images of PGQDs and NGQDs are shown in **Fig. S2**. According to the AFM height profiles, the synthesized materials could be categorized as few-layer GQDs, 1.5–3 nm in height. The AFM image of NGQDs (**Fig. S2** bottom) also shows that they tend to aggregate with an increased height of around 5 nm. GGQDs were synthesized according to the work of Hassan et al.^30^ and were used without characterization.

**Fig. 2.**
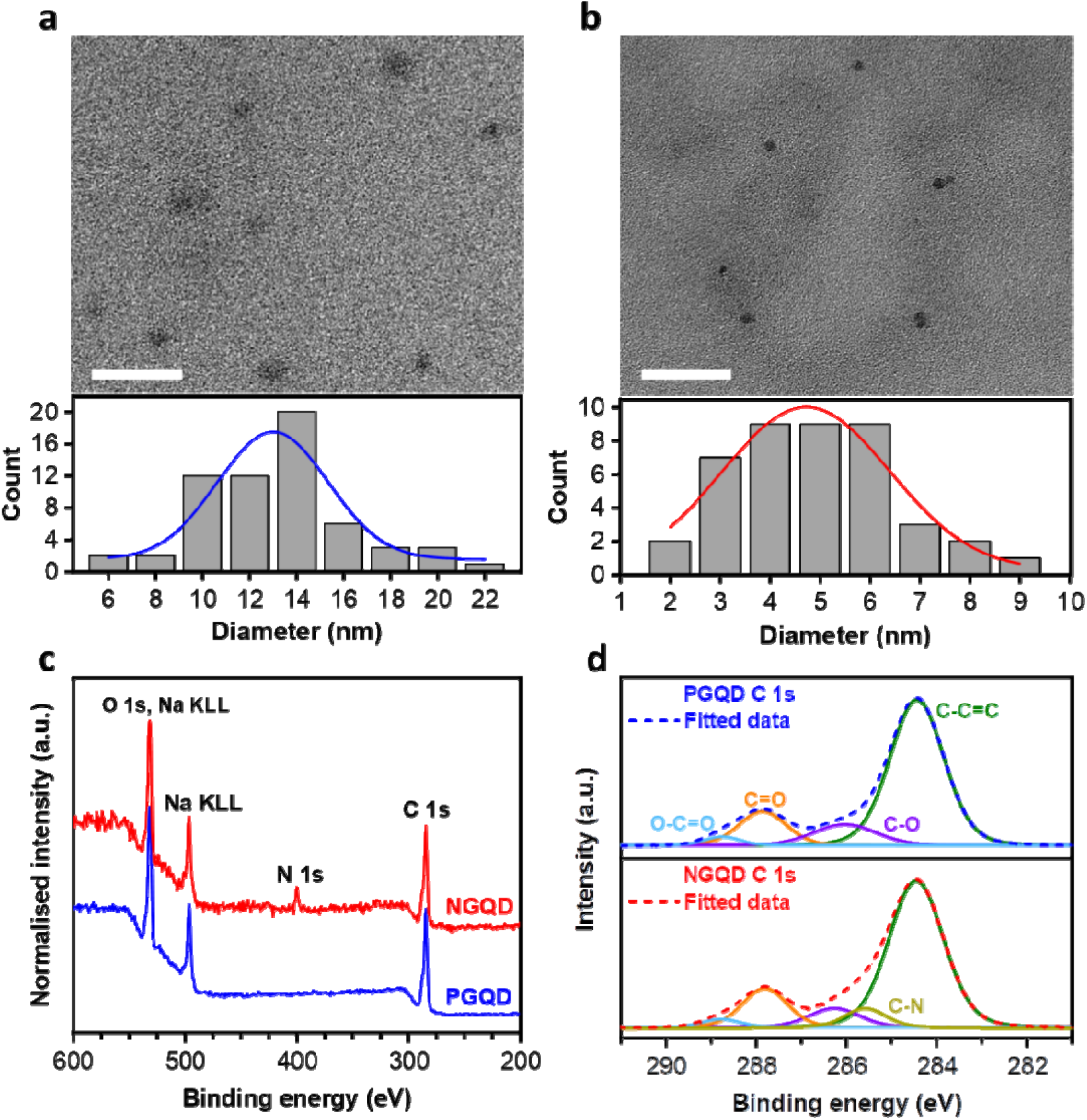
Characterization of GQD samples. TEM images (top) and diameter size distribution (bottom) of PGQDs (a) and NGQDs (b). Scale bars are 50 nm. (c) XPS survey spectra of the as-prepared samples confirming the presence of nitrogen in the NGQD sample. (d) Deconvolution treatment of the C1s spectra of PGQDs and NGQDs (the color of peaks for different bonds are the same for top and bottom panels).

We conducted XPS measurements to analyze the composition of the as-produced GQDs. The XPS survey spectra of the PGQDs and NGQDs exhibited a predominant graphitic C 1s peak at about 284.5 eV and an O 1s peak at about 532 eV (**Fig. 2c**). The O/C atomic ratio for PGQDs and NGQDs was 64.26% and 68.50%, respectively, demonstrating that the NGQDs had a higher level of oxidation than PGQDs, likely due to the shrinkage of PGQDs (**Table S1**). However, the Na content of 5.83% and 5.14% for PGQDs and NGQDs, respectively, and carboxyl group content of 2.75% and 2.46% for PGQDs and NGQDs, respectively, confirmed the reduction of carboxylic acid groups of PGQD by hydrazine (**Table S1**). In addition, a pronounced N 1s peak was observed for the resultant NGQDs, whereas no N signal was detected for the PGQDs (**Fig. 2c**), demonstrating the successful incorporation of N atoms into NGQDs by hydrazine treatment of PGQDs. The N/C atomic ratio was 12.53% (**Table S1**), which is remarkably higher than the NGQDs reported previously.^31^ High-resolution C 1s spectra of PGQD and NGQD are presented in **Fig. 2d**, deconvoluted to their components, including the covalent bonds between C atoms and O and N atoms (contributions in % are given in **Table S1**).

The UV-vis and PL spectra of the synthesized GQDs are provided in **Fig. S3**. The optical absorption edge was 460, 635, and 565 nm, revealing 2.69, 1.95, and 2.19 eV optical bandgaps of PGQDs, NGQDs, and GGQDs, respectively. When excited at 375 nm, all the synthesized GQDs emitted light in the 450–470 nm range. Moreover, the prepared GQDs showed an up-conversion fluorescence in the 400– 500 nm region.

As this work aimed to modulate the photoluminescence of SWCNTs, GQDs were added to an aqueous solution of SWCNTs (containing at least 12 different SWCNT chiralities, HiPCO) suspended with a random 40-nucleotide (nt) ssDNA sequence, called ssDNA-SWCNTs hereafter unless otherwise stated.

The microscopic characterization of ssDNA-SWCNTs and their mixtures with NGQDs and PGQDs are depicted in **Fig. 3**. The STEM image of pure ssDNA-SWCNTs shows long nanotubes with a diameter of around 4 nm. A more precise diameter evaluation is provided by the AFM image showing nanotubes with ∼1 nm diameter (height), in good agreement with the values reported for chiralities in the HiPCO mixture.^32^ The STEM and AFM images of ssDNA-SWCNT+PGQD (**Fig. 3**) reveal a cover of GQDs on the nanotube surface, increasing the height by ∼1.5 nm, which matches the height of pure PGQDs (1.5 nm) represented in **Fig. S2**. As expected, a thicker shell of NGQDs (compared to that of PGQD) is visible around the nanotubes in STEM and AFM images. According to the AFM height profile, the diameter of ssDNA-SWCNT+NGQD conjugates increased to around 6 nm (inset of AFM image of ssDNA-SWCNT+NGQD) at some points. However, it is worth noting that the microscopy images were taken in the solid-state form, and the morphology could be different in the solution phase.

**Fig. 3.**
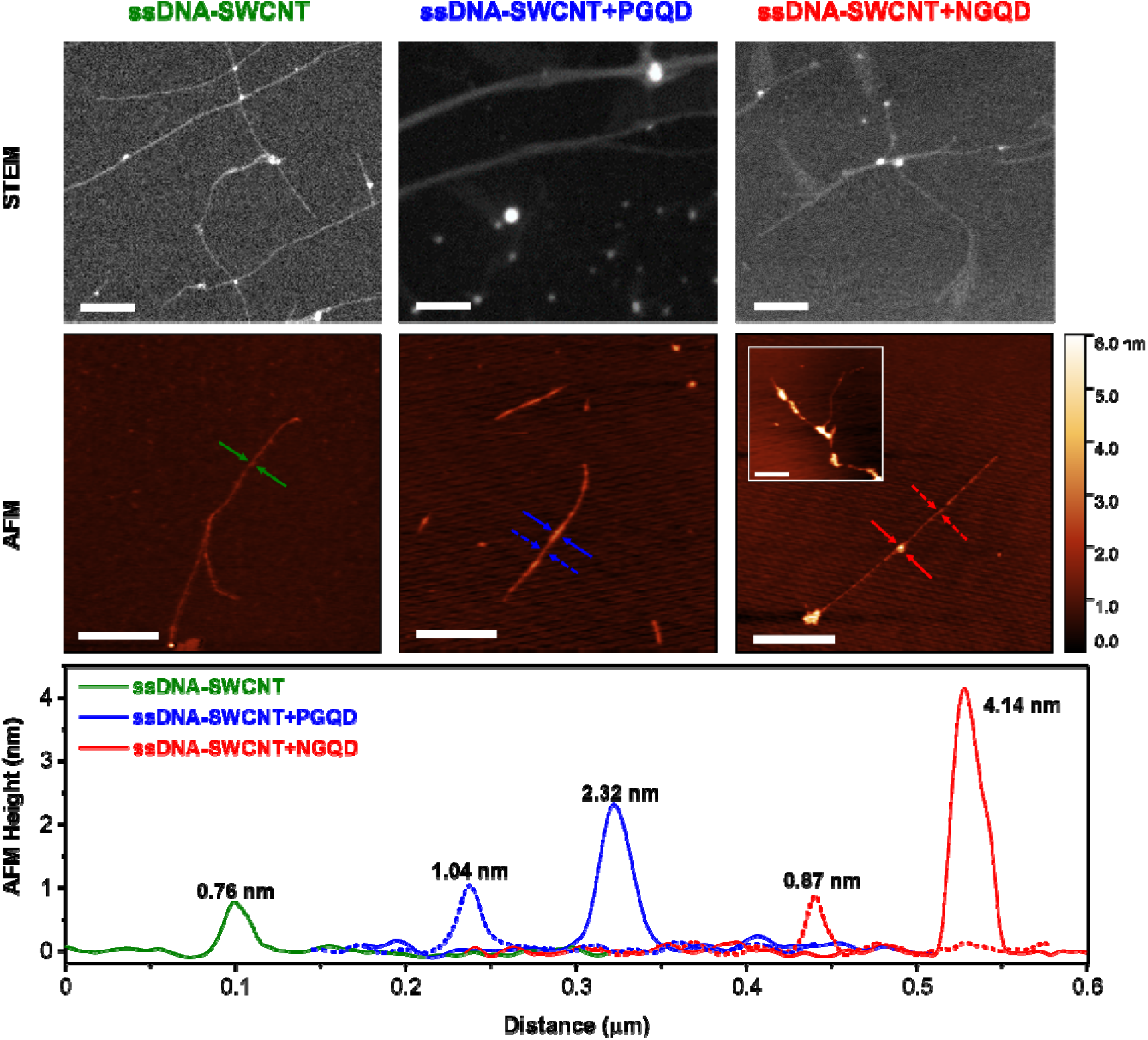
Microscopy characterization of ssDNA-SWCNTs and their mixtures with PGQDs and NGQDs. Top: Representative high-angle annular dark-field STEM images (scale bars are 100 nm). Middle: AFM images (scale bars are 300 nm). Bottom: Height profiles extracted from the corresponding region indicated with arrows in the AFM images. ssDNA-SWCNT is indicated in green, blue represents ssDNA-SWCNT+PGQD, and red shows the ssDNA-SWCNT+NGQD profile.

The absorbance spectrum of ssDNA-SWCNTs (**Fig. S3**) shows different peaks in the 400 – 1300 nm range corresponding to different SWCNT chiralities in the HiPCO mixture. The peaks appearing below 550 nm are mainly attributed to metallic nanotubes, whereas peaks in the 600–900 nm and 900–1300 nm regions are assigned to the E_22_ and E_11_ optical transitions of semiconducting nanotubes, respectively. The absorbance spectra of the ssDNA-SWCNTs in the presence of GQDs were nearly identical to the sum of those for pristine ssDNA-SWCNTs and GQDs solutions, except for the E_11_ transitions of semiconducting SWCNT species. Slight red-shifting along with increases in the peak-to-valley ratio of the peaks was observed above 1100 nm, reflecting a change in aggregation. Nanotube aggregation broadens the resonant absorption peaks and leads to background growth through increased spectral congestion.^33^ The increase in the peak-to-valley ratio was more significant in the presence of NGQDs (red) compared with the PGQDs (blue) and GGQDs (yellow), implying better solubility of ssDNA-SWCNTs when NGQDs are added.

**Fig. 4a** compares the NIR PL excitation-emission (PLE) maps for a solution of ssDNA-SWCNTs before and after adding GQDs. The figure illustrates dramatic increases in the fluorescence intensity of ssDNA-SWCNTs in the presence of PGQDs and NGQDs, both reaching similar intensities after 3 h of incubation. PL enhancement was also achieved upon adding GGQDs to ssDNA-SWCNTs (**Error! Reference source not found**.). However, GGQDs, the control N-doped-GQD sample, were not as effective as NGQDs. As the negative control, the left-side panel in **Fig. 4a** shows that the synthesized PGQDs did not fluoresce in the NIR region while excited in the 500–800 nm range. The same results were attained for NGQDs and GGQDs (**Fig. S4**).

**Fig. 4.**
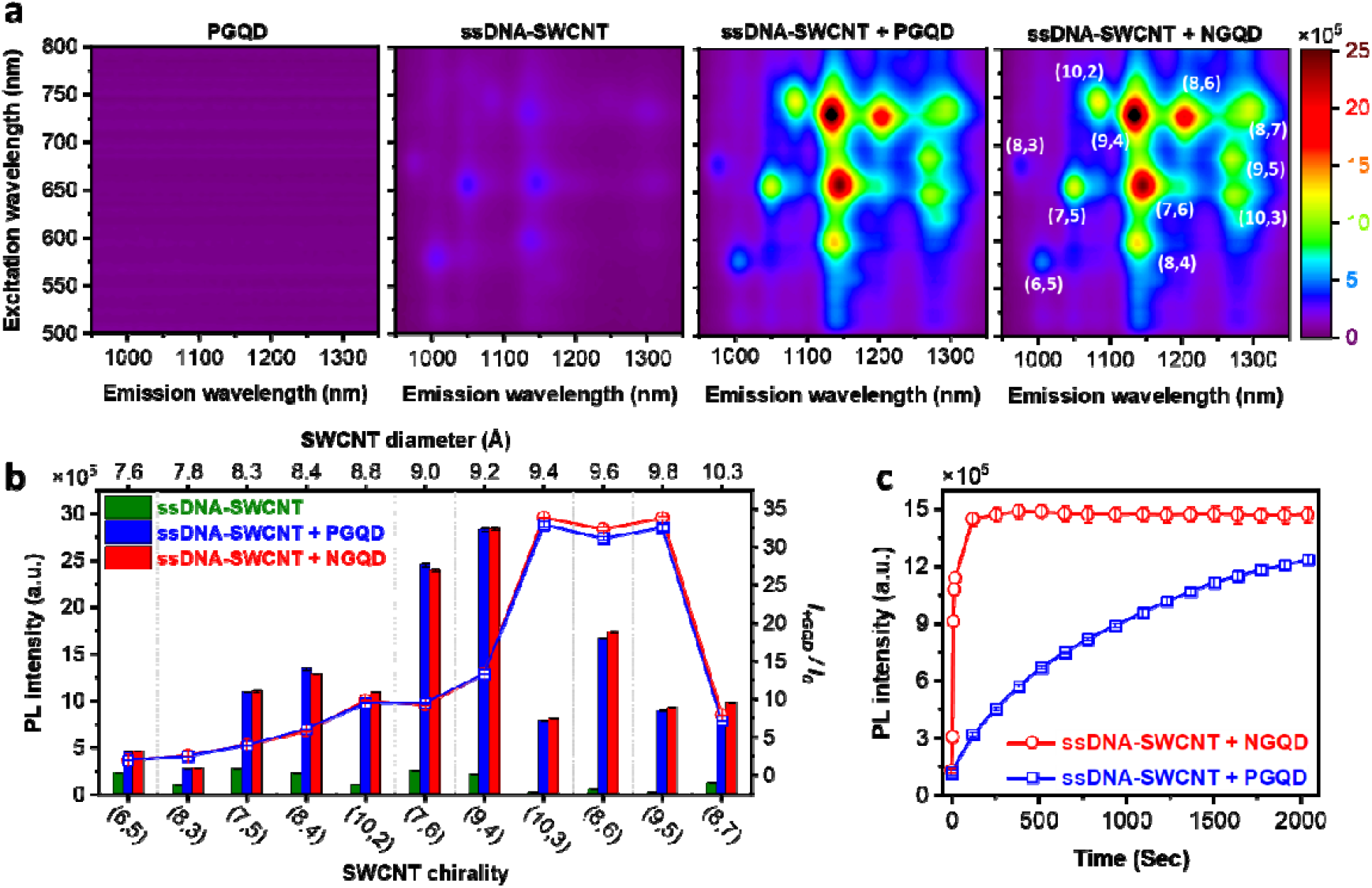
Fluorescence response of ssDNA-SWCNTs to GQDs. (a) NIR-PLE maps of PGQDs (which is similar to that of NGQDs (**Fig. S4**)) and ssDNA-SWCNT (2 mg/L) before and after the addition of PGQDs or NGQDs (with final concentrations of 0.5 mg/ml) following 3 h incubation. (b) PL brightening effect of GQDs for different SWCNT chiralities; absolute PL intensities (bars and left-side axis) along with the brightening factor (scatter-line plot and right-side axis). (c) Kinetic measurements of the influence of GQDs on the fluorescence of ssDNA-SWCNTs. In panel c, PL of (9,4) chirality (excitation at 730 nm and emission at 1135 nm) is presented.

We noted that the fluorescence enhancement when GQDs were added depended on the SWCNT chirality. As depicted in **Fig. 4b**, the increase in the fluorescence intensity was more significant for chiralities of larger diameters. The chiralities with a bright initial state, including (6,5), (7,5), (8,4), (7,6), and (9,4), showed fluorescence intensity increases of up to 2.0, 4.0, 5.8, 9.3, 13.4 times, respectively. This fluorescence enhancement was much more significant for low-QY, larger-diameter (10,3), (8,6), and (9,5) chiralities. They were brightened more than 30 times upon the addition of GQDs. However, an exception was the largest chirality (8,7) in the investigated PLE region; its PL intensity was enhanced by 7.9 times, probably due to a difference in DNA conformation on its surface compared with the lower diameter chiralities. Further investigations through molecular dynamics simulations could help to explore if there are any abrupt changes in the DNA wrapping in this case. A quantitative comparison of the fluorescence changes for different chiralities is presented in **Table S2**.

The kinetics of fluorescence change following the addition of GQDs was also investigated. As shown in **Fig. 4c**, the fluorescence intensity reached a maximum within ∼2 min of adding NGQDs. In the case of PGQD addition, although the fluorescence intensity did not reach a plateau in the studied time range of ∼0.5 h, it is evident in **Fig. 4b** that the final fluorescence intensities for all chiralities are similar to ssDNA-SWCNT+NGQD after 3 h of incubation.

Salem et al.^18^ reported that the configuration of ssDNA around SWCNT changes with the ionic strength of the solution. They showed that the amount of free DNA in the solution decreased with increased salt concentration and demonstrated that the missing ssDNAs were wrapped onto the SWCNTs surfaces. The cations present in the solution acted as shielding particles and decreased the repulsion forces between the phosphate anions of the DNA backbone.^34^ Possessing a large number of carboxylate ions with Na counter cations, GQD can change the ionic strength of aqueous solutions. Therefore, we investigated the influence of GQD addition on the amount of free ssDNA in ssDNA-SWCNT solutions. We found that free ssDNA decreases upon adding GQDs, regardless of their type (**Fig. S5a**). This finding confirms that ssDNA configuration around SWCNTs changes, and more free ssDNA can adsorb onto the nanotube surface.^18^ In addition, we acquired the PL spectra of ssDNA-SWCNTs (with or without GQDs) in several NaCl concentrations ranging from 0 to 2.0 M (**Fig. S5 b** and **c**). As shown in **Fig. S5b**, we observed an increase in the PL intensity of ssDNA-SWCNTs and ssDNA-SWCNTs+PGQDs with increasing salt concentration until around 1.0 M, while that of ssDNA-SWCNTs+NGQDs did not significantly change in this range for it had already reached a maximum (saturated PL) due to the presence of NGQDs. In [NaCl] > 1.0 M, ssDNA-SWCNTs begin to precipitate (visible to the naked eye), probably due to the limited solubility of DNA in high [NaCl],^34,35^ and the PL intensity decreases. From this observation, we conclude that the major role of GQDs in the PL enhancement of ssDNA-SWCNTs is altering the DNA conformation by adjusting the ionic strength.

Recently, Hou et al.^36^ suggested that the PL brightening response upon the addition of reductants is due to a non-localized process of surfactant reorganization around SWCNTs. Studies of isolated nanotubes over a range of aqueous surfactant-SWCNT suspensions indicate that changes in the local dielectric environment and the surfactant coverage of the SWCNTs are essential factors responsible for PL enhancement.^12^ To investigate the effect of GQDs on the surface coverage of SWCNTs, we examined the PL response of surfactant-suspended nanotubes.

As a model surfactant system, we chose SC, which shows a concentration-dependent surface coverage of solubilized particles.^12^ For SWCNTs in particular, Gillen et al.^12^ recently demonstrated that adding neurotransmitters (dopamine and serotonin) to SC-SWCNTs alters the fluorescence emission of the SWCNT. The lower the surface coverage (lower SC concentration), the higher the fluorescence quenching and the larger the peak shifting upon the addition of neurotransmitters. Notably, major fluorescence responses were observed when the SC concentration was below its CMC (∼0.4% w/w in water). Taking this study into account, the PL response of SC-SWCNTs solutions to the addition of GQDS, DTT as a reductant, and potassium ferricyanide (K_3_[Fe(CN)_6_]) as an oxidant were investigated. It was confirmed that ferricyanide ions could be irreversibly adsorbed on the surface of SWCNTs and partially withdrew electrons from the nanotubes, leading to quenching of the nanotubes’ fluorescence.^36^

Expectedly, significant changes in the fluorescence of low surface coverage SC-SWCNTs ([SC] < CMC_SC_) in terms of both intensity and position of the (9,4) peak were recorded upon adding K_3_[Fe(CN)_6_] (**Fig. 5 a** and **b**). An 85% decrease in the fluorescence intensity and 5.2 nm red-shifting in the peak position of SC-SWCNTs were observed at [SC] = 0.02% w/w. In comparison, only 7% decrease in the fluorescence intensity and a slight blue-shift of 0.5 nm in the peak position were detected at [SC] = 0.5% w/w, confirming the surface coverage dependency of SC-SWCNTs. On the other hand, when DTT, a reducing agent, was added to the SC-SWCNT solutions, slight fluorescence changes (quenching and red-shifting) were observed in low surface coverages. In the high surface coverage regime ([SC] > CMC_SC_), though, DTT showed a different behavior compared with the oxidant. Higher fluorescence quenching and slight red-shift in the peak position were recorded for higher SC concentrations. This finding agrees with Hou et al., who explained that SC molecules bind very tightly to SWCNTs, and DTT changes the SC organization around SWCNTs by weakening the SC-SWCNT attraction forces. A similar behavior as DTT was observed in the case of NGQD addition, suggesting this compound’s oxidative nature. PGQDs and GGQDs showed a similar surface coverage dependency as K_3_[Fe(CN)_6_] when added to the SC-SWCNTs series. However, this behavior is most likely related to the direct interaction of GQDs with the SWCNT surface in low surface coverage.

**Fig. 5.**
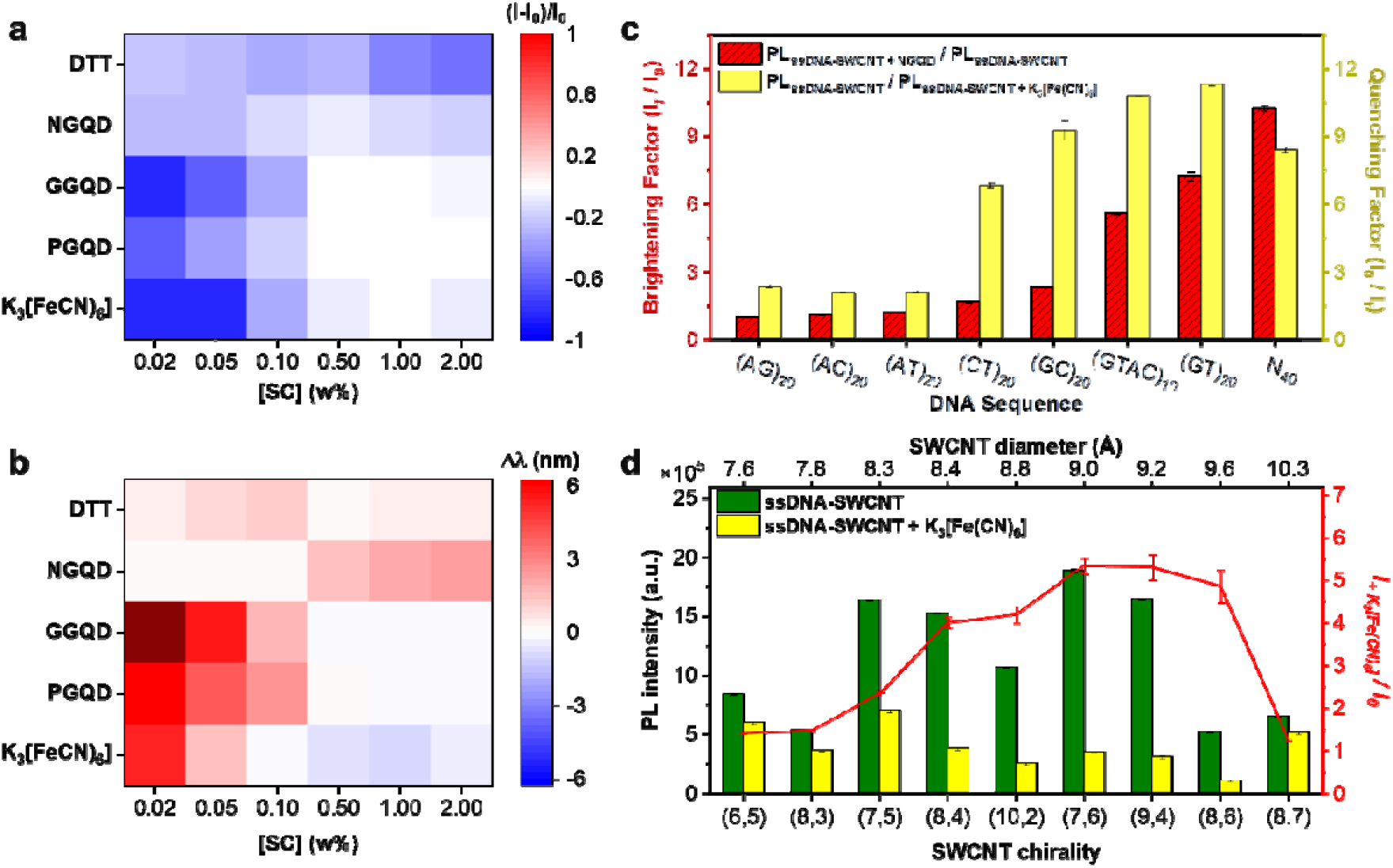
Surface coverage measurements of SC and ssDNA suspended SWCNTs. Effect of several additives, including GQDs, on the fluorescence intensity (a) and peak position (b) of SC-SWCNTs of different SC concentrations. In the heatmaps, red indicates an intensity increase or red-shift in the wavelength position of the (9,4) peak, and blue represents an intensity decrease or blueshift in the wavelength position. **Table S3** summarizes the magnitude of all responses. (c) Comparison of the brightening effect upon the addition of NGQDs and quenching effect upon the addition of K_3_[Fe(CN)_6_] to (9,4)-SWCNT suspended with different 40-nt long ssDNA sequences. (d) PL quenching effect of K_3_[Fe(CN)_6_] for different chiralities of SWCNTs suspended by N40.

As another example of surfactant-suspended SWCNTs, SDS was used to suspend nanotubes. Like DNA-wrapped SWCNTs, SDS-suspended SWCNTs exhibited significant PL brightening upon adding GQDs (**Fig. S6**). Hou and Krauss^14^ found that reductants enhance the PL QY by increasing the radiative decay rate and decreasing nonradiative decay processes when SWCNTs were wrapped by ssDNA or SDS. In contrast to SC, they suggested that the binding between surfactant molecules and nanotubes increases upon adding DTT, which modifies the local dielectric environment in a more nonpolar fashion. Electrolyte-induced brightening of SDS-suspended SWCNTs was also reported by Duque et al..^19^ It was suggested that reorientation of the surfactant partially removes pockets of water from the nanotube surface, playing a crucial role in nonradiative exciton recombination processes. Hence, considering GQDs either as reductants or multi-electrolytes (with Na/C ratio of 10%), they could reorganize the surfactant wrapping, resulting in a PL brightening of SWCNTs.

As shown above, K_3_[Fe(CN)_6_], a small oxidizing agent, could change the fluorescence of SC-SWCNTs in a surface coverage-dependent manner. We used this dependency to measure the surface coverage of ssDNA-SWCNTs qualitatively. Eight ssDNA sequences (40-nt) were used to suspend SWCNTs by the exchange method. Similar to surfactant-SWCNTs, the stronger the adsorption of ssDNA on the nanotube surface, the greater the surface coverage and hence the lower the response to the quenching agent. For instance, it was proved that (AT)_n_ (n = 5, 10, 15, …) sequences were tightly adsorbed on the SWCNT surface, and the resulting sensors only showed responses to highly quenching additives like nitric oxide.^7^ In agreement, the results in **Fig. 5c** show that SWCNTs suspended by adenine-containing ssDNA sequences, including (AT)_20_ and (AT)_15_ (**Fig. S7a**), experience less change in the PL intensity upon the addition of K_3_[Fe(CN)_6_]. On the other hand, It was demonstrated that guanine-containing (high G-content) ssDNAs were lightly adsorbed on the SWCNTs’ surface, resulting in low surface coverage. For example, (GT)_15_ has been used (solely or as an anchor sequence) to suspend SWCNTs because of its great responsivity to different analytes.^37^ Similarly, here, we also observed that (GT)_20_ (**Fig. 5c)** and (GT)_15_ (**Fig. S7b**) were amongst the most responsive sequences against K_3_[Fe(CN)_6_] in the studied ss-DNAs. The response of the identical ss-DNA sequences to the addition of NGQDs was also investigated. As illustrated in **Fig. 5c**, the responsivity order to NGQDs (brightening) was almost the same as that to K_3_[Fe(CN)_6_] (quenching); that is, the sensors with lower surface coverages experienced a greater rise in PL QY and vice versa. Nevertheless, the sensor with random ssDNA (N_40_, containing an equal number of either A, G, C, and T bases) did not match the trend, probably due to the unknown sequence, which could have some particular interactions with K_3_[Fe(CN)_6_] or NGQDs. Of course, the PL intensity of N_40_-wrapped SWCNTs reached near the same level as those wrapped by (AGTC)_10_ (with similar base composition) after the addition of K_3_[Fe(CN)_6_] or NGQDs, and only differed in their initial PL intensities (**Fig. S7a** and **Fig. S8a**).

Moreover, we investigated the influence of the length of wrapping ssDNA on the quenching/brightening effect upon adding K_3_[Fe(CN)_6_]/NGQDs. As (GT)_20_ and the random N_40_ DNA sequences showed the most significant responses, we used (GT)_x_ (x = 5, 10, 15, 20, and 25) and N_y_ (y = 10, 20, 30, 40, and 50), to suspend SWCNTs (panels b and c of Figs. S7 and S8). No meaningful correlation was found between the length of ssDNA and the change in SWCNT fluorescence.

We further examined the quenching effect of ferricyanide on the PL of different SWCNT chiralities suspended by N_40_. As shown in **Fig. 5d**, the quenching magnitude of SWCNT fluorescence increased with their diameter, suggesting lower surface coverage for larger diameters. Roxbury et al.^38^ reported such chirality dependence on ssDNA wrapping around SWCNTs. Using molecular dynamics simulations as well as the PL data for (TAT)_4_-SWCNT, they concluded that (TAT)_4_efficiently suspends small diameter chiralities (e.g. (6,5)) due to interstrand and intrastrand self-stitching hydrogen bonding. However, on larger diameter SWCNTs like (8,7), ssDNA forms a much more disordered and looser structure. Therefore, we can conclude that the surface coverage of larger-diameter SWCNTs suspended by ssDNA is less than that of the small diameters; hence, ferricyanide has more access to the surface of larger-diameter SWCNTs to quench them. Comparing **Fig. 5d** and **Fig. 4c**, we conclude that GQDs brighten larger-diameter SWCNTs more because of their lower surface coverage. Furthermore, in **Fig. 5d**, we observe a drop in the quenching magnitude of the PL of (8,7), similar to what we saw in its PL brightening by GQDs (**Fig. 4c**). This implies the higher surface coverage of (8,7) than that of the smaller-diameter chiralities like (8,6) and (9,4).

In **Fig. 4a** (far left), we demonstrate that GQDs did not fluoresce in the studied NIR region (950–1350 nm) when excited in the 500–800 nm region. However, as shown in **Fig. 6a**, GQDs emitted visible light when excited within the abovementioned region. PGQDs showed a maximum emission peak at around 465 nm when excited at 750 nm. The maximum emission of NGQDs was about 445 nm when excited in the 655-720 nm range. This up-conversion fluorescence is a known phenomenon in GQDs, where the fluorophore emits at shorter wavelengths than the excitation wavelength.^33^ When comparing **Fig. 6a** with **Fig. S3** (right panels), we note that the maximum emission wavelength for the up-converted fluorescence of both PGQDs and NGQDs was quite similar to their normal fluorescence (A = 375 nm). As evident in the right-side panels of **Fig. 6a**, NGQDs were quenched upon adding ssDNA-SWCNTs, while PGQDs were slightly brightened. Since the emission of NGQDs is not in the range where HiPCO chiralities are resonantly excited (above 500 nm), no energy transfer is expected in the mixtures. However, another reason for NGQD quenching in a mixture with ssDNA-SWCNTs could be electron transfer from NGQDs to SWCNTs.

**Fig. 6.**
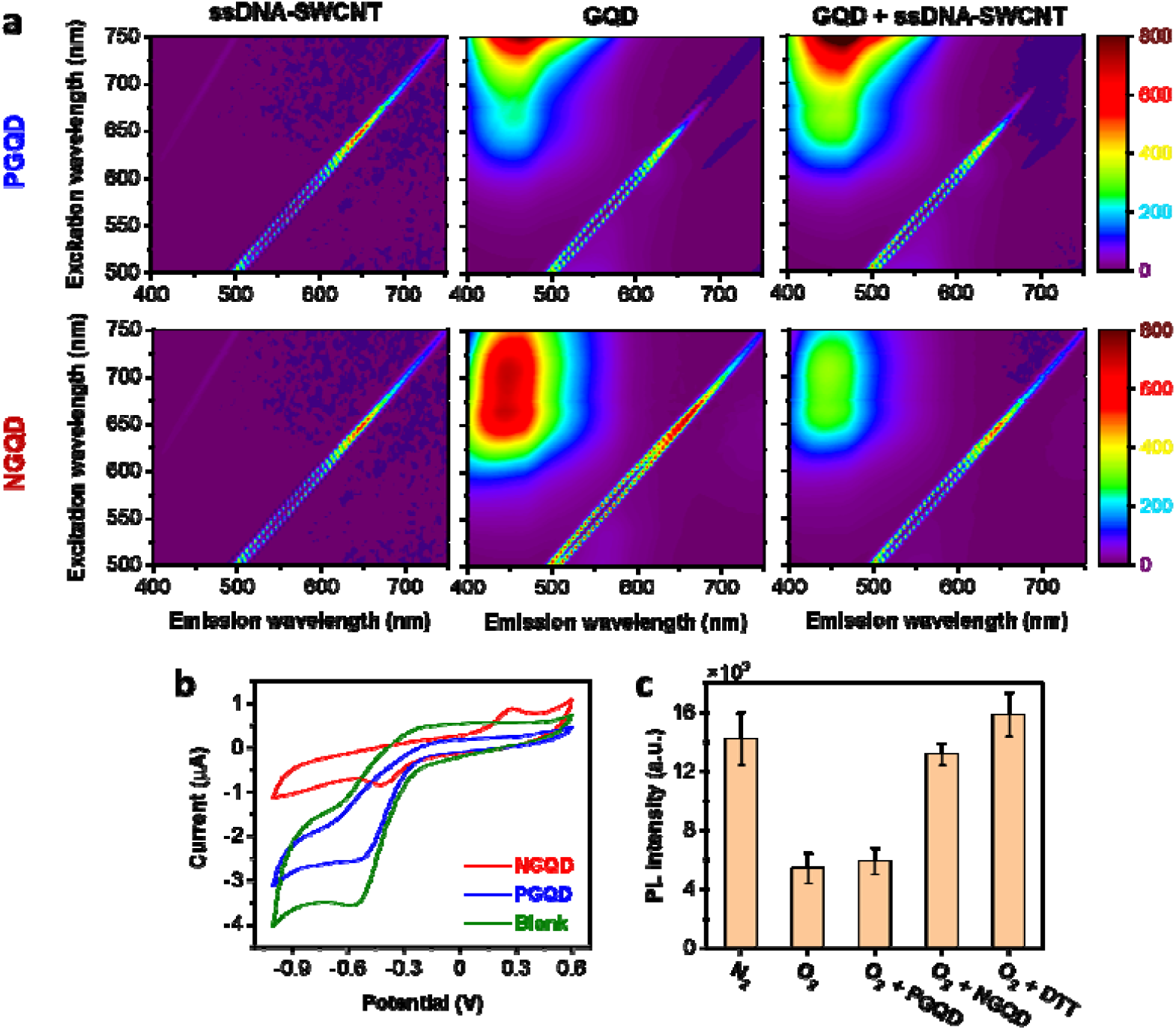
Oxidative behavior of NGQDs for fast brightening of SWCNTs. (a)Visible PLE plots of ssDNA-SWCNT (left side), GQDs (middle), and GQDs+ssDNA-SWCNT conjugates (right side) for PGQD (top) and NGQD (bottom). (b) Cyclic voltammograms of glassy carbon electrode (with a scan rate of 5 mV/s) recorded in NaCl solution (100 mM, blank) and supplemented with GQDs (1 mg/ml). (c) PL response of gas-purged ssDNA-SWCNT solutions before and after adding GQDs (or DTT). PL spectra were collected 15 min after the addition of GQDs, and DTT was used as a control PL intensity of (9,4) chirality (excitation at 730 nm and emission at 1135 nm) is presented.

To investigate if GQDs could transfer electrons, we performed cyclic voltammetry (CV) when their solution was interfaced with a glassy carbon electrode. The cyclic voltammogram of GQDs in 100 mM NaCl solution is shown in **Fig. 6b**. When the potential was swept from –1.0 to +0.6 V, NGQDs produced two redox peaks, with the oxidation one centered at around +0.28 V, with an onset potential of about +0.12 V. CV curves of PGQDs and GGQDs (**Fig. 6b** and **Fig. S9 a-d**) were also investigated in NaCl solution. Neither PGQDs nor GGQDs produced oxidation signals in the studied potential range of –1.0 to +0.6 V. The unique anodic properties of NGQDs are attributed to the introduced hydrazide groups (**Fig. S9e**). It has been reported that refluxing GQDs with hydrazine forms hydrazide groups at the edges of GQDs.^31^ As shown in **Fig. S9e**, the hydrazide group of NGQDs could be chemically oxidized by O_2_ in basic^39^ (and neutral^40^) solutions. All our experiments were carried out at neutral pH, rendering this kind of oxidation likely. Consequently, NGQDs could undergo an electron transfer in a solution of SWCNTs, acting as a reducing agent.

Reducing agents such as DTT, Trolox, and β-mercaptoethanol increase the quantum efficiency of ssDNA-SWCNTs through a transient reduction of defect sites on the SWCNT sidewall.^15^ In the work of Lee et al., the fluorescence intensity of ssDNA-SWCNT solution was enhanced four-fold upon the addition of DTT.^15^ To confirm the rise in fluorescence caused by reduction, they deliberately quenched a solution of ssDNA-SWCNTs by adding an oxidant, i.e., methyl viologen. The fluorescence intensity recovered significantly after adding a reductant, Trolox. In another work, Salem et al. showed that adding an antioxidant (ascorbic acid) to ssDNA-SWCNT solution induces PL brightening.^18^ They also found a similar turn-on response through O_2_ degassing. Nevertheless, the removal of the reducing agent or re-adsorption of oxygen upon exposure to air would decrease the fluorescence intensity of ssDNA-SWCNT solutions to pre-brightening levels. Although ssDNA-SWCNT solution quenches in low ionic strength and acidic conditions, adding reductants to or removing dissolved oxygen from ssDNA-SWCNT solution increases the PL intensity to the same extent. Considering this discussion, we can conclude that NGQDs increase the quantum efficiency of ssDNA-SWCNTs by acting as reducing agents and decreasing oxygen accessibility to the SWCNT surface.

We also performed a series of gas experiments to confirm the reducing function of NGQDs (**Fig. 6c** and **Fig. S10**). Nitrogen-purged and DTT-added samples showed the highest PL intensities with a slight difference due to the incomplete desorption of O_2_ when degassing (by nitrogen purging). Expectedly, NGQDs increased the PL intensity of the oxygen-purged ssDNA-SWCNT solution. As explained above, NGQDs were oxidized **(Fig. S9e**) in the presence of O_2_ (and consumed it), thereby quenching SWCNTs. Also, a slight increase in the PL intensity was observed upon PGQD addition. Therefore, both PGQDs and NGQDs could increase the QY of ssDNA-SWCNT solutions but through different mechanisms. It seems that NGQDs kinetically speed up the brightening of SWCNTs. To reach a thermodynamically stable bright state, both NGQDs and PGQDs change the nanotubes’ environment (e.g., by reconfiguring ssDNA around SWCNT).

The applicability of PL brightening of ssDNA-SWCNTs by adding GQDs was investigated through two kinds of experiments. First, increasing PL QY of individual SWCNTs was observed through NIR confocal microscopy. Although recent developments in NIR confocal microscopy enable increased single-molecule imaging resolution, the improved resolution occurs at the expense of significant fluorescence losses. From the confocal NIR images shown in **Fig. 7a**, we observed that ssDNA-SWCNTs complexes experienced a significant PL enhancement in the presence of NGQDs. It was also revealed that after extensive washing (3 times) of the ssDNA-SWCNTs+NGQD mixture (conducted to remove NGQDs), the PL intensity of individual ssDNA-SWCNTs was much more than their intensities before the addition of NGQDs. This is a unique behavior of NGQDs, as removing reductants like DTT (used as brightening agents) from the ssDNA-SWCNTs solution leads to complete quenching of the fluorescence of SWCNTs to their original state, even in wide-field NIR microscopy.^15^ To confirm that NGQDs were entirely removed in the washing process, the visible fluorescence image of the respective samples was recorded (**Fig. 7b**). The washed sample (SWCNTs+NGQD; washed) did not show any fluorescence in the visible region. Moreover, due to the electron transfer, the visible fluorescence intensity of the SWCNTs+NGQD sample was lower than that of pure NGQDs.

**Fig. 7.**
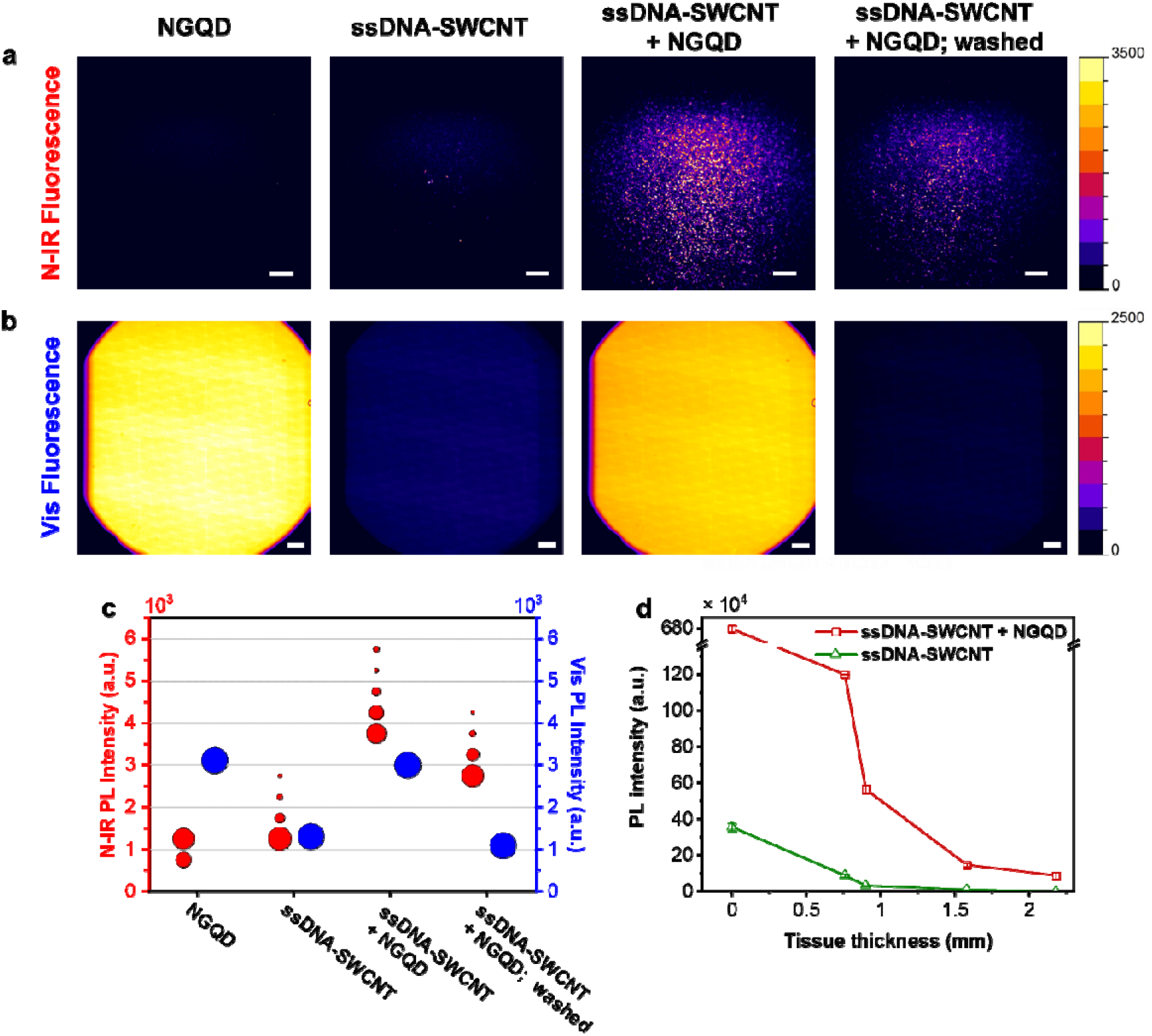
Application of PL brightening of SWCNT by adding GQDs. (a) Confocal NIR microscopy of individual ssDNA-SWCNTs before and after the addition of NGQD. In all images, samples were excited at 640 nm, and emission was collected above 980 nm using a long-pass filter. (b) Visible photoluminescence microscopy of the respective samples in panel a. Samples were excited at 370 nm, and the emission was recorded in the visible region. Scale bars are 10 µm in all panels “a” and “b” images. (c) Frequency distributions of NIR PL (left-side red axis) intensities of individual ssDNA-SWCNTs in the presence and absence NGQDs and the visible PL of the respective samples (right-side blue axis). The 200 brightest 2 × 2 pixelated regions in a and b were considered for the histograms. The surface of circles is proportional to the frequency of the regions with the respective intensities. (d) The penetration depth of PL of ssDNA-SWCNT in ham tissues of different thicknesses in the absence and presence of NGQDs.

Besides their biocompatibility, another advantage of GQDs over reducing agents is their relatively large size. This is important when making implantable (in-vivo) sensors with fine membranes that keep sensing material (i.e., SWCNT) and allow the transfer of analytes. Due to their small size, toxic reducing agents may flow into the tissues and quench the sensor. A quantitative comparison of different samples in terms of the NIR PL intensities of individual SWCNTs and visible PL intensities of NGQDs is illustrated in the circle histograms (**Fig. 7c** and **Table S4**).

To explore the potential application of the ssDNA-SWCNTs for deep tissue sensing, we investigated the penetration depth of ssDNA-SWCNTs in ham tissue. A dramatic decrease in the PL intensity of ssDNA-SWCNTs ((9,4) peak) with increasing tissue thickness was observed (**Fig. 7d**). No signal was recorded for thicknesses more than 1 mm. After the addition of NGQDs, the fluorescence signal was significantly enhanced. Though the PL intensity decreased with tissue depth, an appreciable signal was recorded even for thicknesses more than 2 mm.

## Conclusion

This work achieved bright ssDNA-SWCNT fluorescent sensors through conjugation with GQDs. The impacts of pristine medium-level oxidation GQDs (PGQDs; fabricated via a straightforward route of pyrolysis of citric acid) and NGQDs (product of the reaction of hydrazine with PGQD) on the PL of ssDNA-SWCNT were compared. Thermodynamically, quite the same brightening factors were obtained through their conjugation with PGQDs and NGQDs for all chiralities after 3 h. The amount of free ssDNA in the ssDNA-SWCNT solution decreased upon adding both GQDs. This confirms that ssDNA configuration around SWCNTs changes, and more free DNA can adsorb onto the nanotube surface.^18^ This configurational change could alter the SWCNT surface coverage and hence change the fluorescence. To prove this, we qualitatively measured the surface coverage of ssDNA-SWCNT of different ssDNA wrappings by adding a ferricyanide solution. The results confirmed that low surface coverage SWCNTs, with (GT)_20_ for example, experience a significant increase in brightness while high surface coverage SWCNTs, with (AT)_20_ for example, have a minor change. This is a crucial point that must be considered in designing sensors. (GT)_y_ sequences have shown great responsivity to different analytes.^6^ However, researchers usually use high concentrations of (GT)_y_-SWCNTs that are good enough to be detected with NIR cameras. Such high detection limits represent a weakness of such sensors. We can overcome this drawback by using GQDs to sensitize low-concentration (GT)_y_-SWCNTs fluorescence sensors. Our in-vitro assessment of very low concentration SWCNTs (2 mg/L) wrapped by (GT)_20_ in deep tissues via single-molecule NIR confocal microscopy revealed substantial increases in the confocal microscopy signal (2.87-fold) and penetration depth (>two-folds) of (GT)_20_-SWCNTs upon the addition of NGQDs.

From a kinetic point of view, PL enhancement of ssDNA-SWCNTs by PGQDs and NGQD was different at short durations (< 1 h). With NGQDs, ssDNA-SWCNTs could reach the maximum brightness in a few minutes but did not reach a plateau even after 1 h of adding PGQDs. Compared to the literature, we concluded that NGQDs could speed up the brightening process by passivating the defect sites on the SWCNTs sidewall^15^ as well as neutralizing the quenching effect of dissolved oxygen. Previously, it was confirmed that fluorophores undergo a quenching process through admixture with SWCNTs due to a photoinduced excited electron transfer from the fluorophore to the SWCNT.^21^ The same effect in the case of NGQDs was observed in this work. Also, cyclic voltammetry measurements demonstrated an oxidation peak for NGQDs, which was not detected for PGQDs. In addition, it was reported that similar nitrogen-doped GQDs react with oxygen and release nitrogen in solution^31^, which can induce ssDNA-SWCNT fluorescence.

To conclude, this study demonstrates a simple approach for modulating and sensitizing ssDNA-SWCNTs’ fluorescence. Sensors with enhanced fluorescence and increased *in-vivo* imaging depths can be prepared using low-cost, biocompatible GQDs that are easy to synthesize and handle. In addition, due to their highly efficient photoluminescence, GQDs have been used in various sensing applications. Therefore, mixing with SWCNTs enables one to create bi-modal fluorescence sensors active in both visible and NIR light regions.

## Acknowledgements

The authors are thankful for support from the Swiss National Science Foundation Project No. 200021_184822.

