## Supplementary material for "Photoluminescence brightening of single-walled carbon nanotubes through conjugation with graphene quantum dots": SI

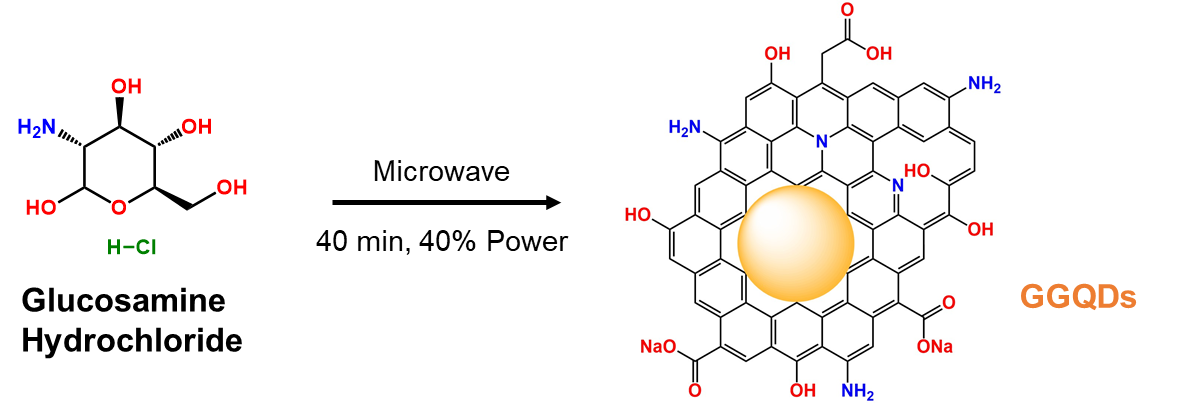

**Fig. S1** Synthesis of GGQDs through microwave heating of glucosamine‐HCl aqueous solution.

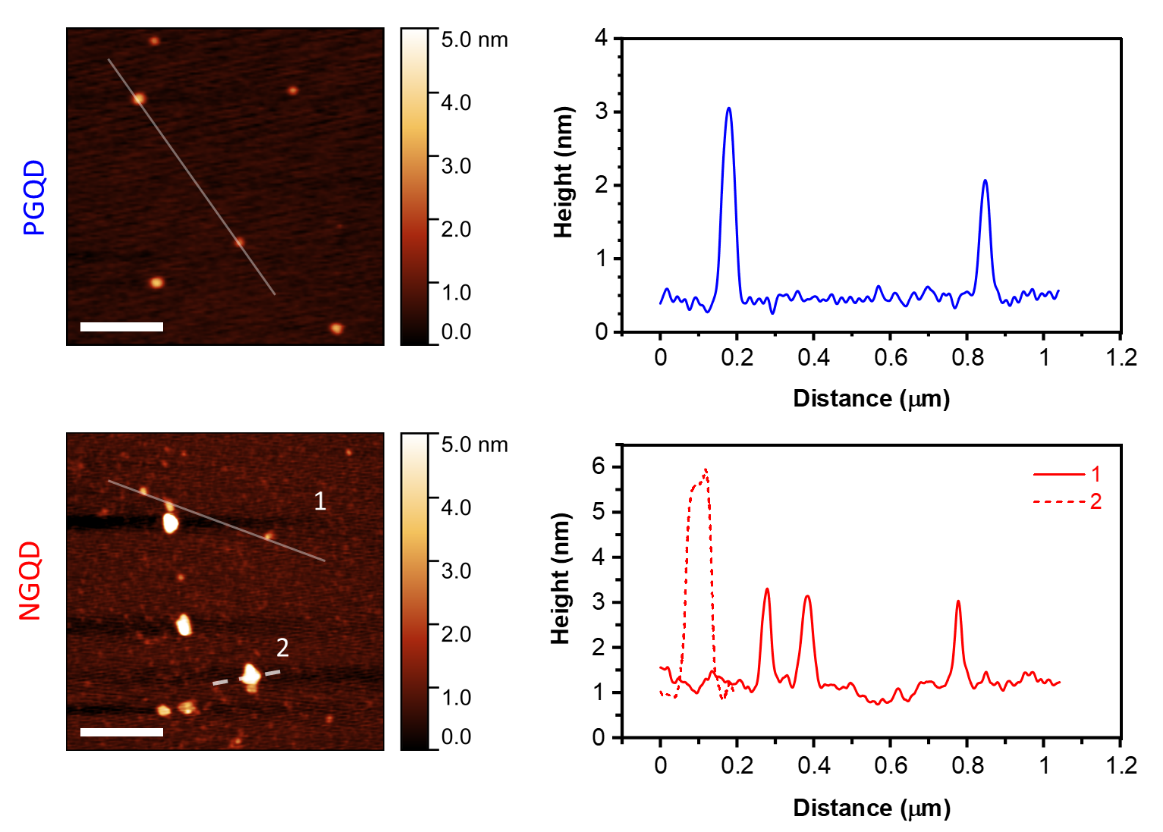

**Fig. S2** Atomic force microscopy (AFM) imaging of GQD samples. Representative AFM topographies (left side, scale bars: 300 nm) and height profiles (right side) correspond to the AFM's white lines.

**Table S1** Atomic composition of PGQDs and NGQDs according to XPS analysis

|  | **Atomic % (Survey XPS spectrum)** | | | |  | **Atomic % (C 1s deconvolution)** | | | | |
| --- | --- | --- | --- | --- | --- | --- | --- | --- | --- | --- |
| **Sample** | C | N | O | Na |  | C=C-C  (284.5 eV) | C-N  (285.6 eV) | C-O  (286.3 eV) | C=O  (287.8 eV) | O-C=O  (288.8 eV) |
| **PGQDs** | 57.39 | N/A | 36.88 | 5.83 |  | 70.73 | N/A | 14.54 | 11.97 | 2.75 |
| **NGQDs** | 52.40 | 6.57 | 35.89 | 5.14 |  | 68.28 | 6.43 | 8.07 | 14.76 | 2.46 |

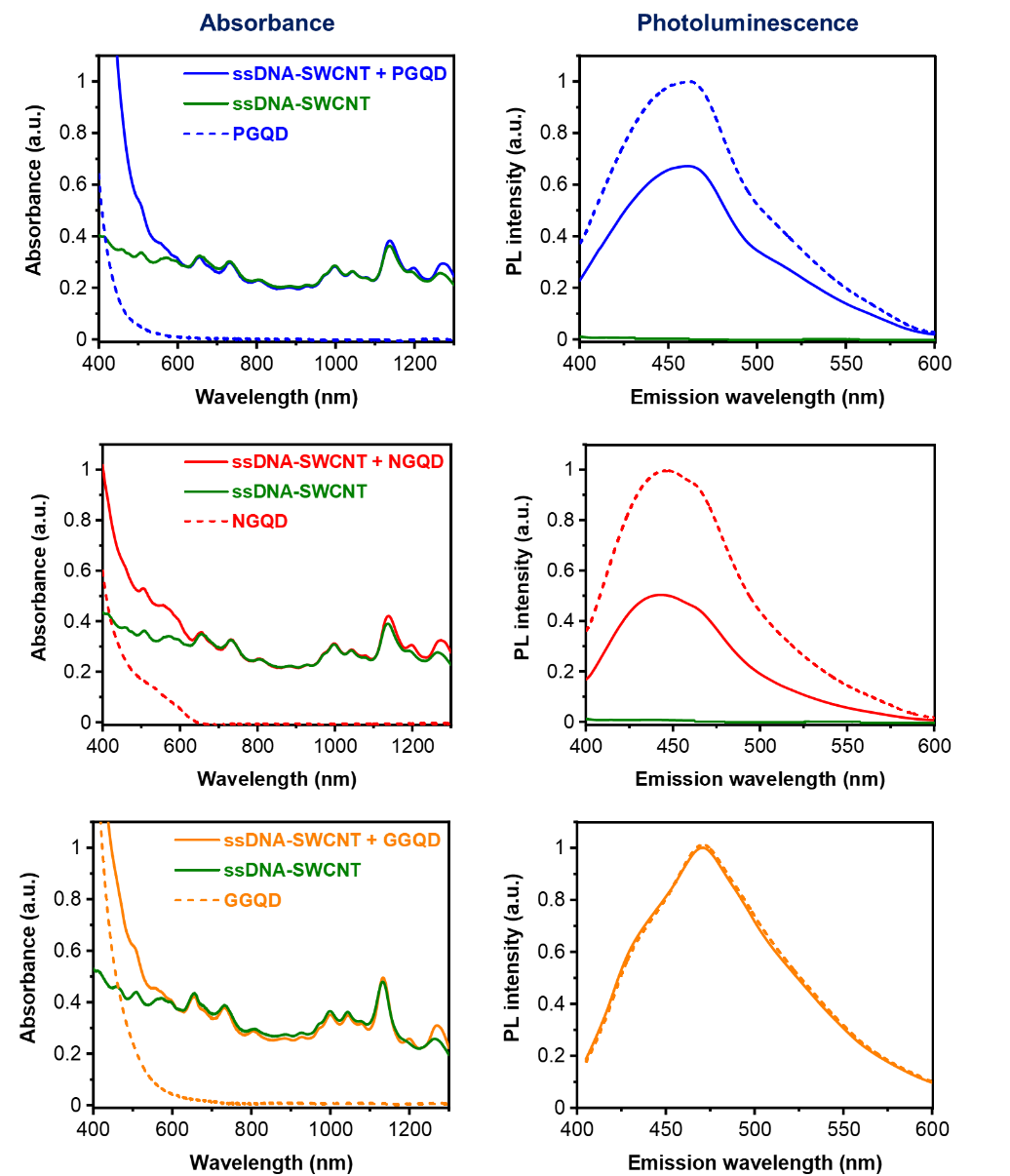

**Fig. S3** Absorbance (left side) and PL spectra (right side, excited at 375 nm) of pure GQDs, ssDNA-SWCNT, and ssDNA-SWCNT + GQD samples. The legends on the right side are the same as those for the corresponding panels on the left side.

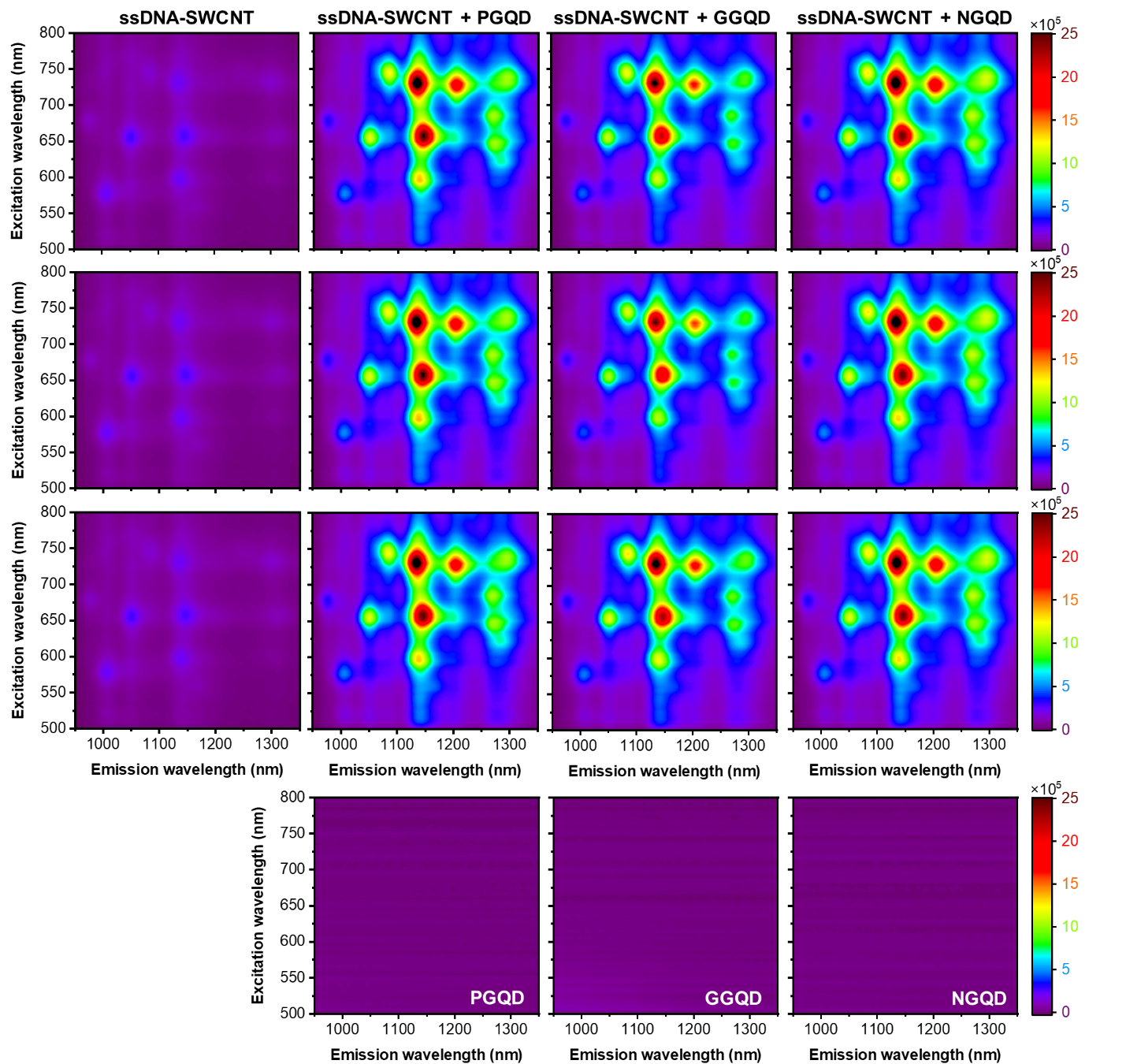

**Figure S4** NIR-PLE maps ( three replicates) of ssDNA-SWCNT (2mg/l) before and after the addition of different GQDs with different functional groups (0.1 mg/ml) following 3 h incubation. NIR-PLE maps of pure GQDs are shown at the bottom raw.

**Table S2** PL brightening effect of NGQDs and PGQDs for different ssDNA-SWCNTs chiralities.

| SWCNT chirality | Diameter ($Å$) |  | PL_ssDNA-SWCNT_  Intensity (a.u.)  (× 10^4^) |  | PL_ssDNA-SWCNT + GQD_ / PL_ssDNA-SWCNT_ | | |
| --- | --- | --- | --- | --- | --- | --- | --- |
|  |  |  |  |  | PGQD | GGQD | NGQD |
| (6,5) | 7.57 |  | 22.99 ± 0.04 |  | 2.03 ± 0.02 | 1.89 ± 0.13 | 2.00 ± 0.01 |
| (8,3) | 7.82 |  | 10.78 ± 0.16 |  | 2.54 ± 0.01 | 2.51 ± 0.17 | 2.63 ± 0.01 |
| (7,5) | 8.29 |  | 27.61 ± 0.11 |  | 3.98 ± 0.02 | 4.21 ± 0.24 | 4.01 ± 0.04 |
| (8,4) | 8.40 |  | 22.11 ± 0.19 |  | 6.08 ± 0.07 | 5.13 ± 0.29 | 5.84 ± 0.03 |
| (10,2) | 8.84 |  | 11.15 ± 0.07 |  | 9.49 ± 0.03 | 9.63 ± 0.47 | 9.85 ± 0.05 |
| (7,6) | 8.95 |  | 25.80 ± 0.11 |  | 9.50 ± 0.08 | 8.28 ± 0.44 | 9.27 ± 0.05 |
| (9,4) | 9.16 |  | 21.20 ± 0.03 |  | 13.34 ± 0.08 | 12.13 ± 0.62 | 13.40 ± 0.07 |
| (10,3) | 9.36 |  | 2.39 ± 0.06 |  | 33.01 ± 0.35 | 33.11 ± 0.13 | 33.92 ± 0.04 |
| (8,6) | 9.66 |  | 5.36 ± 0.05 |  | 31.17 ± 0.23 | 26.62 ± 1.38 | 32.42 ± 0.20 |
| (9,5) | 9.76 |  | 2.74 ± 0.17 |  | 32.60 ± 0.41 | 30.52 ± 1.64 | 33.85 ± 0.21 |
| (8,7) | 10.32 |  | 12.27 ± 0.05 |  | 7.18 ± 0.06 | 6.35 ± 0.29 | 7.96 ± 0.02 |

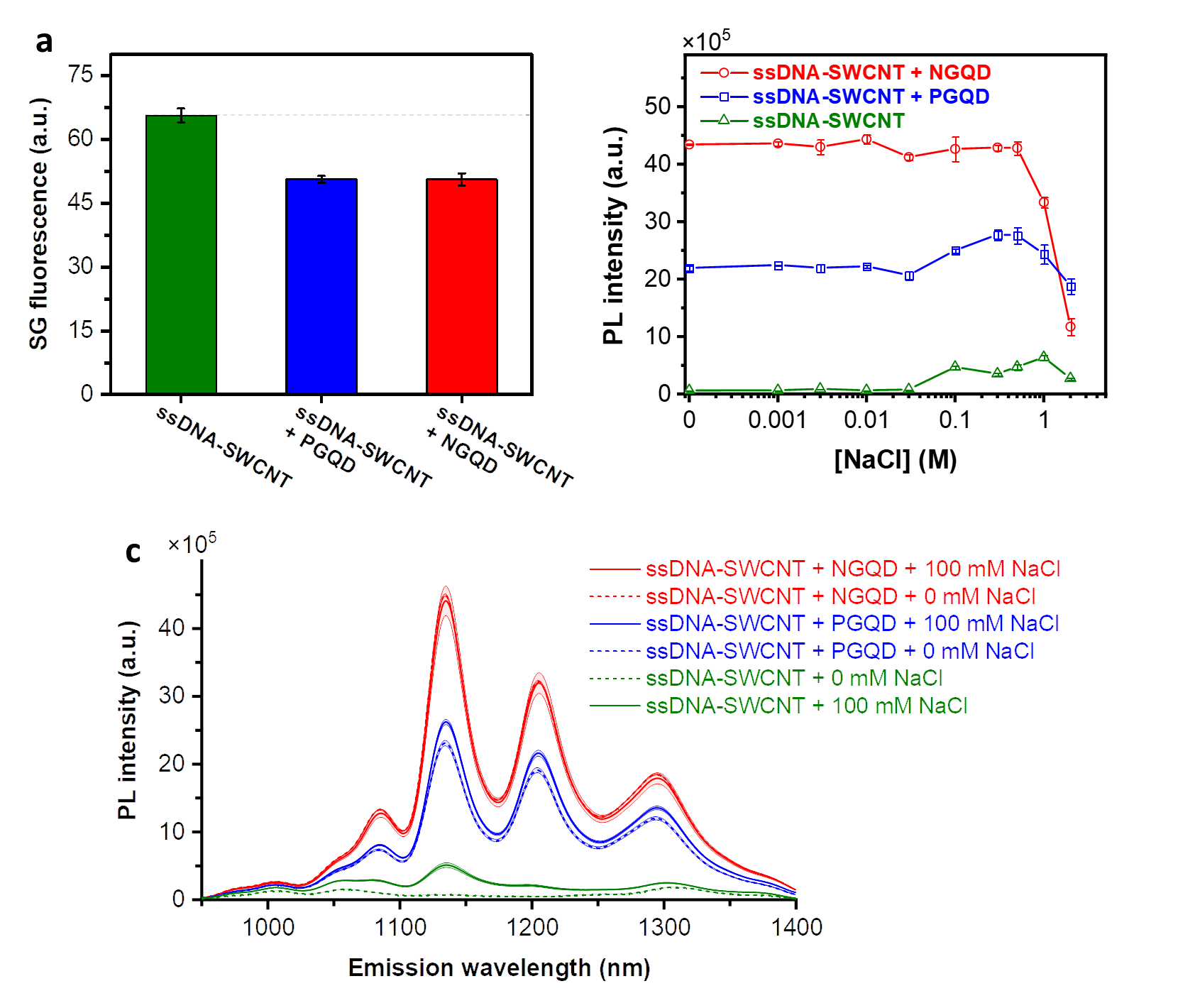

**Fig. S5** Influence of the ionic strength of solution on the ssDNA configuration around nanotubes and PL intensity of ssDNA-SWCNT solutions. (a) Fluorescence of Sybr Gold (SG) showing the amount of free (unbound) ssDNA in ssDNA-SWCNT ((9,4) chirality) solutions before and after the addition of GQDs. (b) Effect of ionic strength (salt (NaCl) concentration) on the PL of ssDNA-SWCNT samples in the absence and presence of PGQDs or NGQDs. (c) PL spectra of the samples (excited at 730 nm) in 0 and 100 mM NaCl solutions in the absence and presence of PGQDs or NGQDs. In the spectra, the central line represents the average spectrum, with the shaded regions representing 1σ standard deviation (of 3 technical replicates).

**Table S3** Intensity change and peak position shifting for the SC-SWCNT solutions of different SC concentrations on adding different additives. This table corresponds to the main text's heatmaps in Fig. 6 a and b.

| [SC]  (W%) |  | *(I-I_0_)/I_0_* | | | | |  | $\Delta\lambda$ (nm) | | | | |
| --- | --- | --- | --- | --- | --- | --- | --- | --- | --- | --- | --- | --- |
|  |  | K_3_[Fe(CN)_6_] | PGQD | GGQD | NGQD | DTT |  | K_3_[Fe(CN)_6_] | PGQD | GGQD | NGQD | DTT |
| 0.02 |  | -0.85 | -0.61 | -0.85 | -0.24 | -0.23 |  | 5.28 | 6.21 | 14.56 | 0.00 | 0.35 |
| 0.05 |  | -0.82 | -0.36 | -0.61 | -0.25 | -0.26 |  | 1.55 | 3.97 | 5.48 | 0.09 | 1.05 |
| 0.1 |  | -0.31 | -0.18 | -0.32 | -0.16 | -0.31 |  | -0.01 | 2.70 | 1.69 | 0.19 | 1.33 |
| 0.5 |  | -0.07 | -0.01 | -0.01 | -0.08 | -0.28 |  | -0.55 | 0.17 | -0.01 | 1.57 | 0.12 |
| 1 |  | 0.00 | 0.00 | 0.00 | -0.15 | -0.47 |  | -0.94 | -0.15 | -0.24 | 1.99 | 0.35 |
| 2 |  | -0.07 | 0.00 | -0.02 | -0.17 | -0.54 |  | -0.33 | -0.02 | -0.03 | 2.24 | 0.42 |

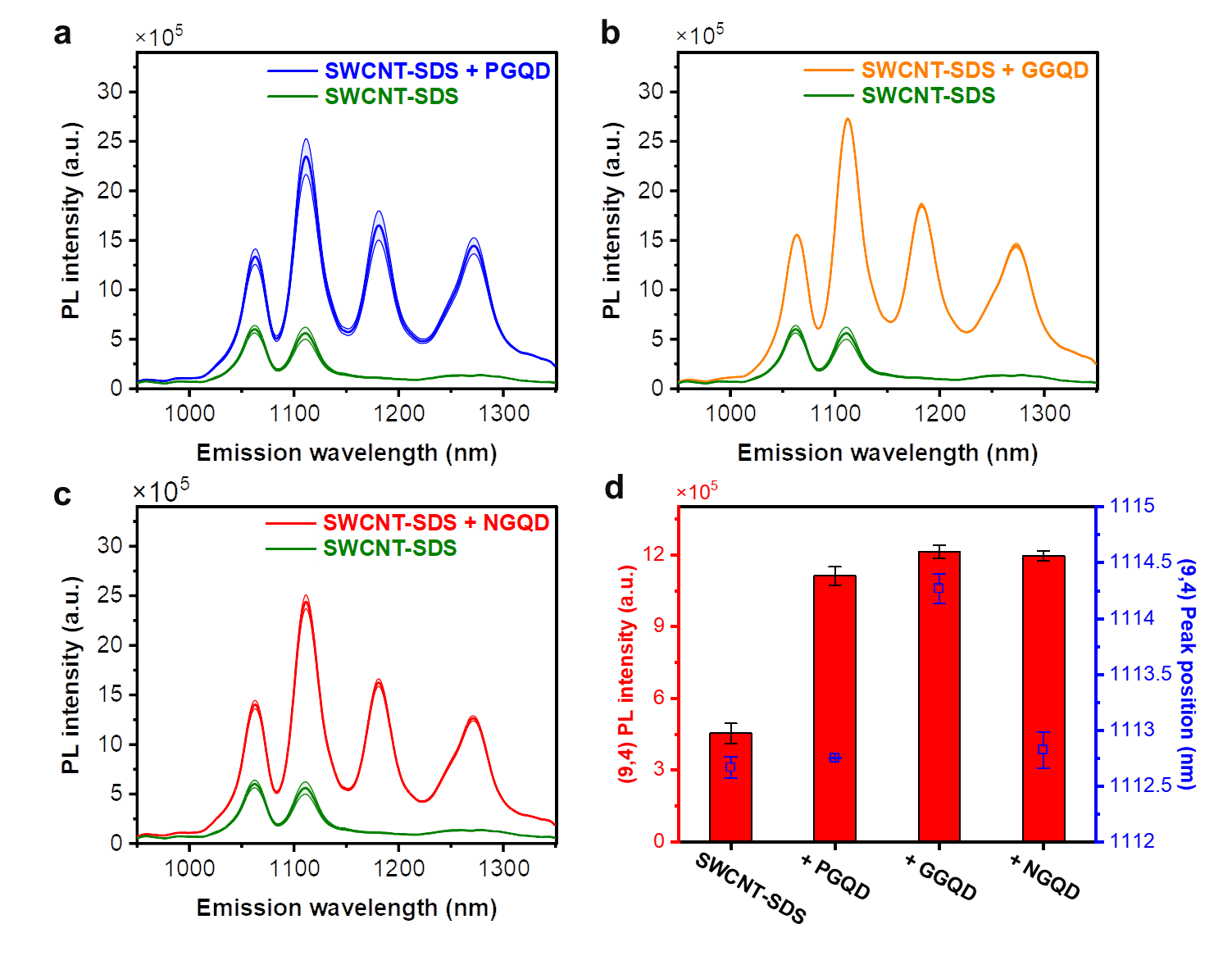

**Fig. S6** PL brightening of SDS-SWCNTs on adding different types of synthesized GQDs. PL spectra of SDS-SWCNTs ([SDS] = 2.0% w/w) before and after the addition of PGQDs (a), GGQDs (b), and NGQDs (c). The spectra were collected after 10 sec of excitation at 730 nm. (d) PL intensity and wavelength for the (9,4) emission peak.

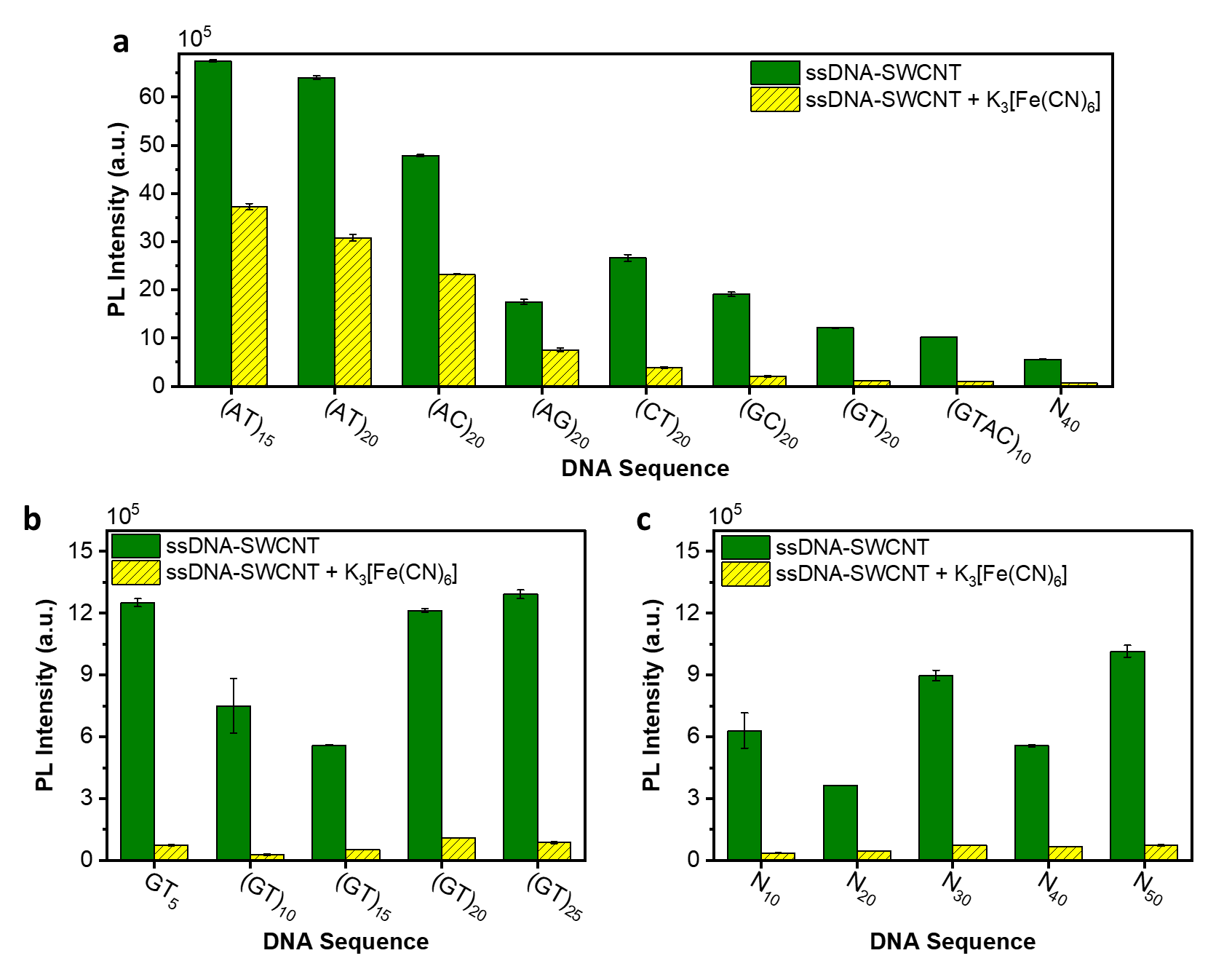

**Fig. S7** PL quenching effect of K_3_[Fe(CN)_6_] for SWCNTs ((9,4) chirality, excited at 730 nm with emission at 1135 nm) suspended by ssDNAs of different sequences and/or lengths. (c) PL intensities of SWCNTs wrapped by different 40-nt long ssDNA sequences before and after the addition of K_3_[Fe(CN)_6_]. (b) PL intensities of SWCNTs wrapped by (GT)_x_ ssDNA sequences before and after the addition of K_3_[Fe(CN)_6_]. (c) PL response of ssDNA-SWCNTs wrapped by random ssDNA sequences of different lengths to the addition of K_3_[Fe(CN)_6_].

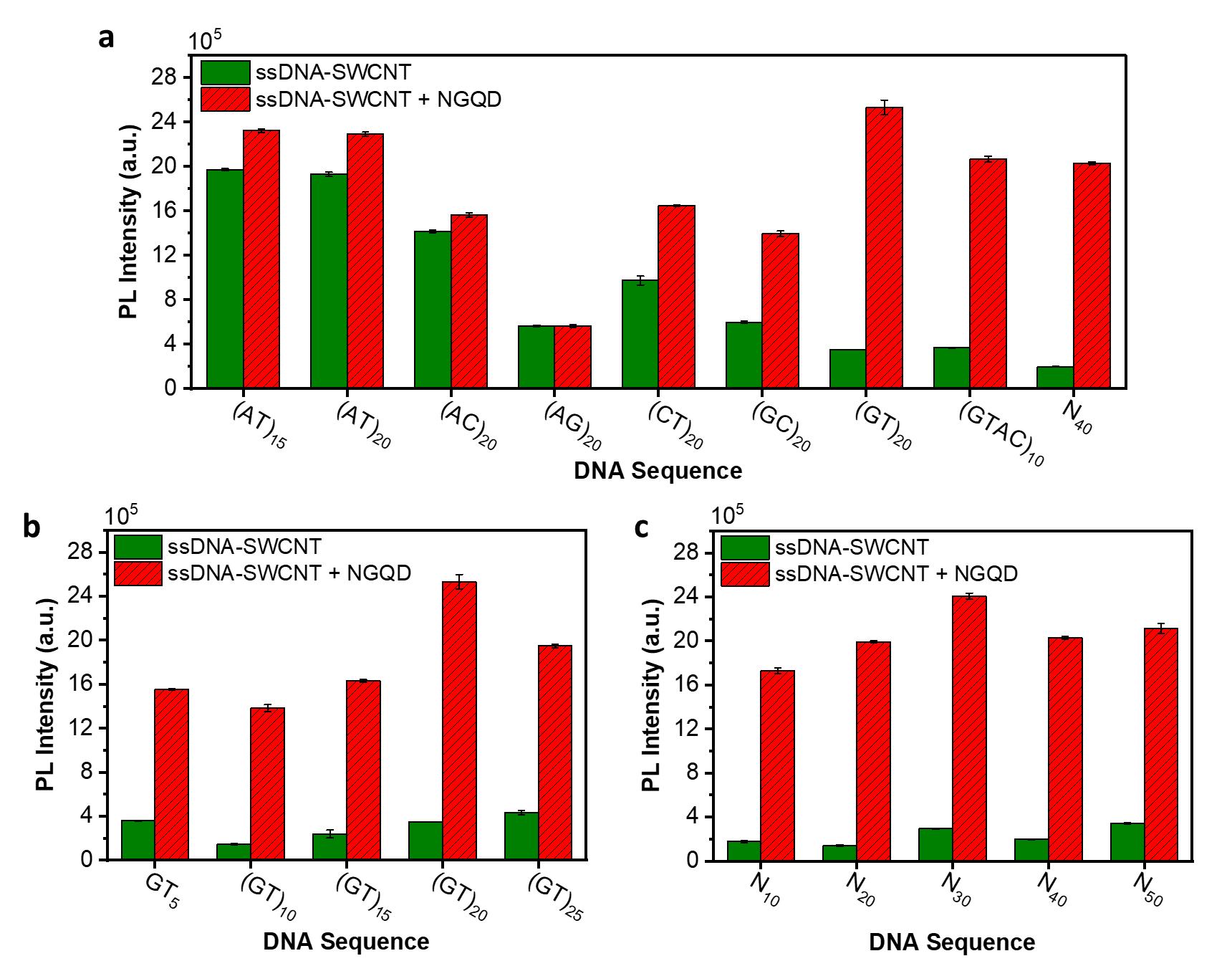

**Fig. S8** PL brightening effect of NGQD for SWCNTs ((9,4) chirality, excited at 730 nm with emission at 1135 nm) suspended with ssDNAs of different sequences and/or lengths. (a) PL intensities of SWCNTs wrapped by different 40-nt long DNA sequences before and after the addition of NGQDs. (b) PL intensities of SWCNTs wrapped by (GT_)x_ DNA sequences before and after adding NGQDs. (c) PL response of SWCNTs wrapped by random ssDNA sequences of different lengths to the addition of NGQDs.

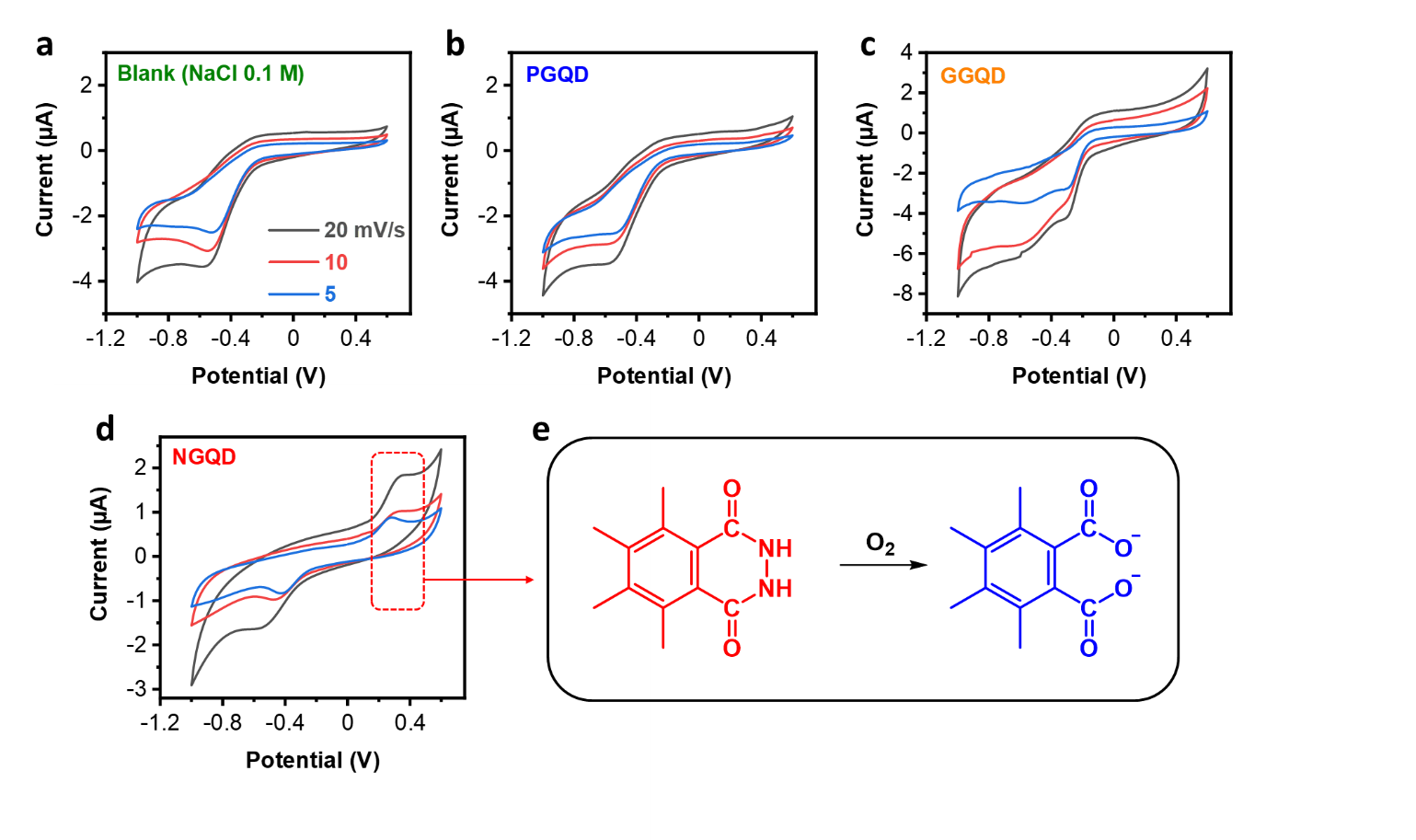

**Fig. S9** Cyclic voltammograms of glassy carbon electrode recorded in (a) buffer (NaCl solution, 0.1 M) or supplemented with (b) PGQDs, (c) GGQDs, and (d) NGQDs at different scan rates (colors for different scan rates are the same in all graphs). (e) Oxidation of edge hydrazide group of NGQDs in the presence of oxygen.

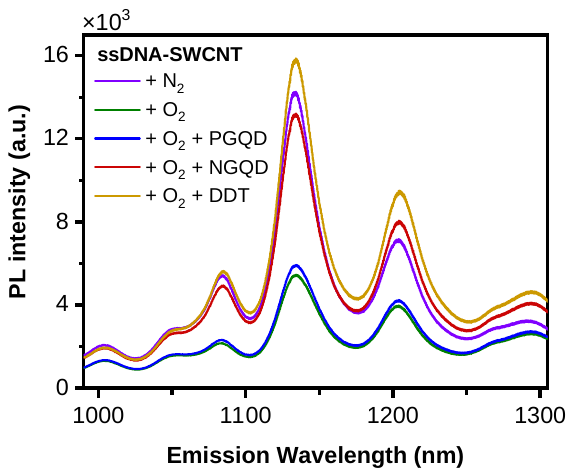

**Fig. S10** PL spectra of the ssDNA-SWCNT (excited at 730 nm) purged with nitrogen or oxygen after adding different additives. This graph corresponds to Fig. 5c in the main text.

**Table S4** Average and standard deviation of intensities of the 200 brightest 2 × 2 pixelated regions in Fig 8 of the main text.

|  | NGQD | ssDNA-SWCNT | ssDNA-SWCNT + NGQD | ssDNA-SWCNT + NGQD; washed |
| --- | --- | --- | --- | --- |
| Mean Visible fluorescence  (Std) | 3319.63  **(**12.07**)** | 1197.17  (4.31) | 3016.07  (11.98) | 1090.15  (4.33) |
| Mean NIR fluorescence  (Std) | 1027.89  (46.54) | 1442.58  (286.65) | 4149.15  (499.53) | 2919.36  (344.97) |
